# Single-cell multi-omic analysis of fibrolamellar carcinoma reveals rewired cell-to-cell communication patterns and unique vulnerabilities

**DOI:** 10.1101/2024.12.11.627911

**Authors:** Alaa R. Farghli, Marina Chan, Marc S. Sherman, Lindsay K. Dickerson, Bo Shui, Manabu Nukaya, Andreas Stephanou, Rosanna K. Ma, Brian J. Pepe-Mooney, Colton J. Smith, Donald Long, Paul R. Munn, Adrian McNairn, Jennifer K. Grenier, Michael Karski, Sean M. Ronnekleiv-Kelly, Venu G. Pillarisetty, Wolfram Goessling, Taranjit S. Gujral, Khashayar Vakili, Praveen Sethupathy

## Abstract

Fibrolamellar carcinoma (FLC) is a rare malignancy disproportionately affecting adolescents and young adults with no standard of care. FLC is characterized by thick stroma, which has long suggested an important role of the tumor microenvironment. Over the past decade, several studies have revealed aberrant markers and pathways in FLC. However, a significant drawback of these efforts is that they were conducted on bulk tumor samples. Consequently, identities and roles of distinct cell types within the tumor milieu, and the patterns of intercellular communication, have yet to be explored. In this study we unveil cell-type specific gene signatures, transcription factor networks, and super-enhancers in FLC using a multi-omics strategy that leverages both single-nucleus ATAC-seq and single-nucleus RNA-seq. We also infer completely rewired cell-to-cell communication patterns in FLC including signaling mediated by SPP1-CD44, MIF-ACKR3, GDF15-TGFBR2, and FGF7-FGFR. Finally, we validate findings with loss-of-function studies in several models including patient tissue slices, identifying vulnerabilities that merit further investigation as candidate therapeutic targets in FLC.

## Introduction

FLC was officially classified by the World Health Organization (WHO) in 2010 as a distinct type of liver cancer characterized by thick intra-tumoral fibrous bands, predominantly affecting adolescents and young adults (*1–6*). There is currently no standard of care. Surgical resection is the only chance for cure; however, this is not a viable option for many patients who present with advanced metastatic disease at the time of diagnosis (*4*). In 2014, the hallmark genetic anomaly of FLC was identified as a ∼400kb somatic, heterozygous deletion leading to an in-frame fusion of exon 1 of *DNAJB1* and exons 2-12 of *PRKACA* (*7*), resulting in the formation of a functional protein chimera (DNAJ-PKAc) that drives tumor initiation in mice (*8*, *9*) and remains relevant for tumor maintenance as well (*10*). The vast majority of FLC patients harbor this chimera (*11*). Targeting and suppressing the chimera is a compelling therapeutic strategy, however past and ongoing efforts have met with significant challenges due in large part to off-target effects on wild-type protein kinase A (PKA) (*12–14*). Therefore, there is a critical need to identify pathways disturbed downstream of DNAJ-PKAc that will reveal molecular dependencies and alternative therapeutic targets (*4*, *15*, *16*).

Over the past decade, several studies have reported genome-scale analyses of FLC tumors (*17– 24*). These include chromatin activity using ChRO-seq (*22*), gene expression through RNA-seq (*11*, *22*), and miRNA expression via small RNA-seq (*23*). These studies uncovered FLC-specific gene markers, enhancer and super enhancer landscapes, and transcriptional and post-transcriptional regulatory networks. Despite these advances, a notable limitation of these studies is their reliance on whole tissue analysis, which obscures cell type specific contributions. The constituent cell types in the tumor microenvironment (TME) and their abundance in FLC relative to non-malignant liver (NML) remain undefined. Moreover, the relevance of each cell type in the TME to the overall molecular landscape of FLC is unknown. This represents an important knowledge gap that merits detailed investigation, especially given that TME and the cell-to-cell communication therein is critical to the development and progression of cancer.

The increased tractability of single-cell genome-scale technologies presents a compelling opportunity to tackle these unresolved questions. Herein, we present the first-ever single-cell study on FLC tumors, encompassing more than 150,000 single cells. We employed both single-nucleus ATAC-seq (snATAC-seq) and single-nucleus RNA-seq (snRNA-seq) to define the constituent cell types of FLC and NML tissue, identify the cell types that are most altered in abundance in FLC, and determine cell type specificity of FLC marker genes (including *SLC16A14*, reported as an FLC marker in 2017 (*20*, *22*)) and miRNAs (including miR-10b, reported as an FLC marker in 2022 (*23*)). We additionally leveraged snATAC-seq (*25*) data to deconvolute the bulk signal from ChRO-seq, and thereby resolve the cell types associated with previously defined FLC-specific super enhancers (including the *SLC16A14*-associated super enhancer, reported as FLC-specific in 2020 (*22*)) and transcriptional networks (including CREB, reported as rewired in FLC in 2020 (*22*)). We also utilized snRNA-seq (*26*) to infer completely new and rewired signaling pathways and cell-to-cell communication patterns in FLC relative to NML. Finally, we provided experimental evidence for specific predictions, most notably the contribution of SLC16A14 to FLC tumor cell viability, with loss-of-function studies in several different models of disease, including patient tumor tissue slices. The study offers the highest-resolution molecular landscape of FLC to date, and the results provide a wealth of new biological insight and potential vulnerabilities that merit further investigation as candidate therapeutic targets in FLC.

## Results

### snATAC-seq and snRNA-seq analysis of human FLC tumors

To decode the tumor biology of FLC at single-cell resolution, we obtained resected frozen human primary tumors alongside adjacent non-malignant liver (NML) tissue from patients. All FLC tumors and NML tissue samples were disassociated and prepped for combinatorial indexing for snATAC-seq or droplet-based snRNA-seq library preparation (Fig. 1A). We compared various quality control (QC) metrics across all samples (fig. S1, S2). After detailed QC analysis, including doublet removal (Table S1&2), we obtained profiles of 32,536 nuclei through snATAC-seq, and 111,133 nuclei through snRNA-seq. We used only primary FLC and NML samples to generate the snATAC-seq and snRNA-seq datasets. Samples used in each assay were derived from different patients.

**Fig. 1.**
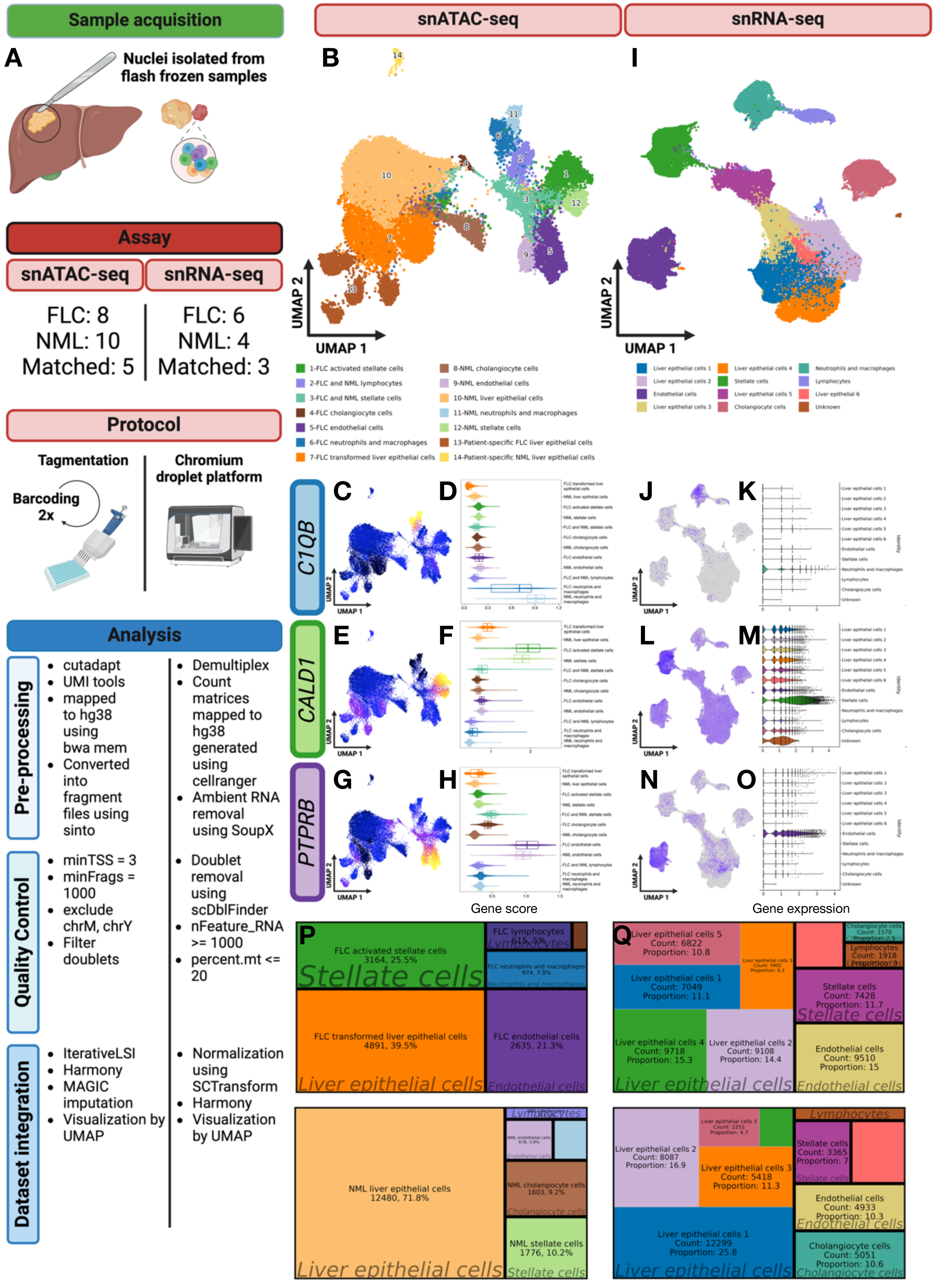
Overview of single-nuclei ATAC-seq and single-nuclei RNA-seq workflows in fibrolamellar carcinoma tumors and non-malignant liver tissue. **(A)** Schematic depicting workflow from sample acquisition to quality control metrics. Liver and protocol cartoon were created using BioRender.com. (B and I) UMAPs of single-nuclei ATAC-seq (snATAC-seq, left; number of nuclei = 32,536) and single-nuclei RNA-seq (snRNA-seq, right; number of nuclei = 111,133), with cells colored by cell type. (C, E, G, J, L, N) Signal for established markers of various liver cell types overlayed atop the snATAC-seq (left; C, E, and G) and snRNA-seq (right; J, L, and N) UMAPs. (D, F, H, K, M, P) Violin plots of ArchR gene activity scores (left; D, F, and H) and Seurat gene expression (right; K, M, and O) for established cell type markers. (P and Q) Treemap showing proportions of cell types depicted in FLC (top row) and NML (bottom row) within snATAC-seq (left) and snRNA-seq (right) datasets. Top panel treemaps correspond to FLC, bottom panel treemaps correspond to NML. UMAP, Uniform Manifold Approximation and Projection; FLC, fibrolamellar carcinoma; NML, non-malignant liver.

The dataset normalization and integration methods are highlighted in Fig. 1A and described further in Materials and Methods. Clusters were visualized for both datasets through uniform manifold approximation and projection (UMAP) (Fig. 1B, I). Patient metadata was also overlayed on the UMAP (snATAC-seq, fig. S3; snRNA-seq, fig. S4). Cells before and after QC are also reported (Supplementary Table 1 and 2). There was no observable stratification based on sex or age and, overall, integration by Harmony substantially removed large batch effects.

However, there may be some minor patient-specific and batch differences that remain unresolved by Harmony integration (*27*) (fig. S3 and S4). Cell type annotations were performed by first identifying enriched markers in each cluster and then overlapping these with known liver cell type markers (Fig. 1C, E, and G). We demonstrated cell type marker specificity using violin plots that depict ArchR calculated gene activity scores (Fig. 1D, F, and H). We then confirmed cell type annotations using an overlap analysis of cluster-enriched genes with previously defined liver cell type markers (*28*) in snATAC-seq (fig. S5A) and snRNA-seq (fig. S5B). The patterns of cell type specificity of different markers were very well-matched between the two datasets (Fig. 1J-O). The clusters for snATAC-seq stratified most cell types based on FLC or NML tissue status (fig. S6A) unlike snRNA-seq, which on the other hand captured more liver epithelial cell subtypes (fig. S7A). These observations likely stem at least in part from the distinct dimensionality reduction techniques employed by ArchR (*29*) (snATAC) and Seurat (*30*) (snRNA). We observed very similar proportions of different cell types in both datasets (Fig. 1F, G, fig. S6B, S7B). Notably, we detected a substantive increase in relative abundance of FLC endothelial and FLC activated stellate cells in both datasets, though particularly striking with snATAC-seq (Fig. 1F).

We then delved deeper into each patient’s contribution to the clusters. In the snATAC-seq dataset, tissue status predominantly determined cluster assignment, with slight variations among patients (fig. S6C). Conversely, patient contributions remained consistent across all clusters in the snRNA-seq dataset (fig. S7C), albeit with a few patient-specific clusters in both datasets. Our snATAC-seq results were further validated using another publicly available analysis pipeline, SnapATAC2 (fig. S8A-G) (*31*). SnapATAC2 identifies the same cell types, calculates similar proportions, and observes comparable cellular contributions and composition from patient samples as ArchR and Seurat. Interestingly, SnapATAC2 does not delineate cell types by tissue status as distinctly as ArchR, possibly due to differences in dimensionality reduction techniques.

### Single nuclei sequencing resolves cell-type specificity of FLC markers and TF networks

A decade of research has yielded numerous FLC tumor markers, but lacking cell type specificity. Both snATAC-seq and snRNA-seq offer an opportunity to bridge this critical knowledge gap.

With snATAC-seq, we visualized the cell types by showing z-score of gene activity scores for various FLC markers (Fig. 2A). Additionally, we leveraged snATAC-seq to visualize activity at microRNA (miRNA) loci, as standard snRNA-seq cannot capture miRNAs. We also visualized patterns of expression across cell types for the same FLC markers using snRNA-seq (Fig. 2B). UMAP visualization clearly shows specificity for FLC gene and miRNA markers across liver epithelial, endothelial, and stellate cells (Fig 2C, E, G, I, K, M). We confirmed cell type specificity of FLC markers using violin plots (Fig 2D, F, H, F, J, L, and N). Notably, in snATAC-seq, we found that miR-10b is primarily sourced from non-epithelial cells in FLCs. This unexpected finding strongly suggests that epithelial-based cell models are insufficient to investigate the function of this miRNA in FLC. A recent study in glioblastoma demonstrated that targeting miR-10b together with another miRNA, miR-21, which is also of interest in FLC (*23*), can regress tumor burden and metastatic potential (*32*), and this merits further investigation in FLC. Using SnapATAC2, we observe similar FLC marker patterns in our snATAC-seq dataset (fig. S8H). Finally, through snRNA-seq, we visualized higher expression of VCAN in FLC activated stellate cells. This observation is significant as VCAN is implicated in extracellular matrix modulation and acts as a pro-fibrotic factor, potentially contributing to the thick fibrotic bands characteristic of FLC. The utilization of single-cell sequencing technology reveals which cellular compartments FLC markers are active within, unveiling potential therapeutic targets.

**Fig. 2.**
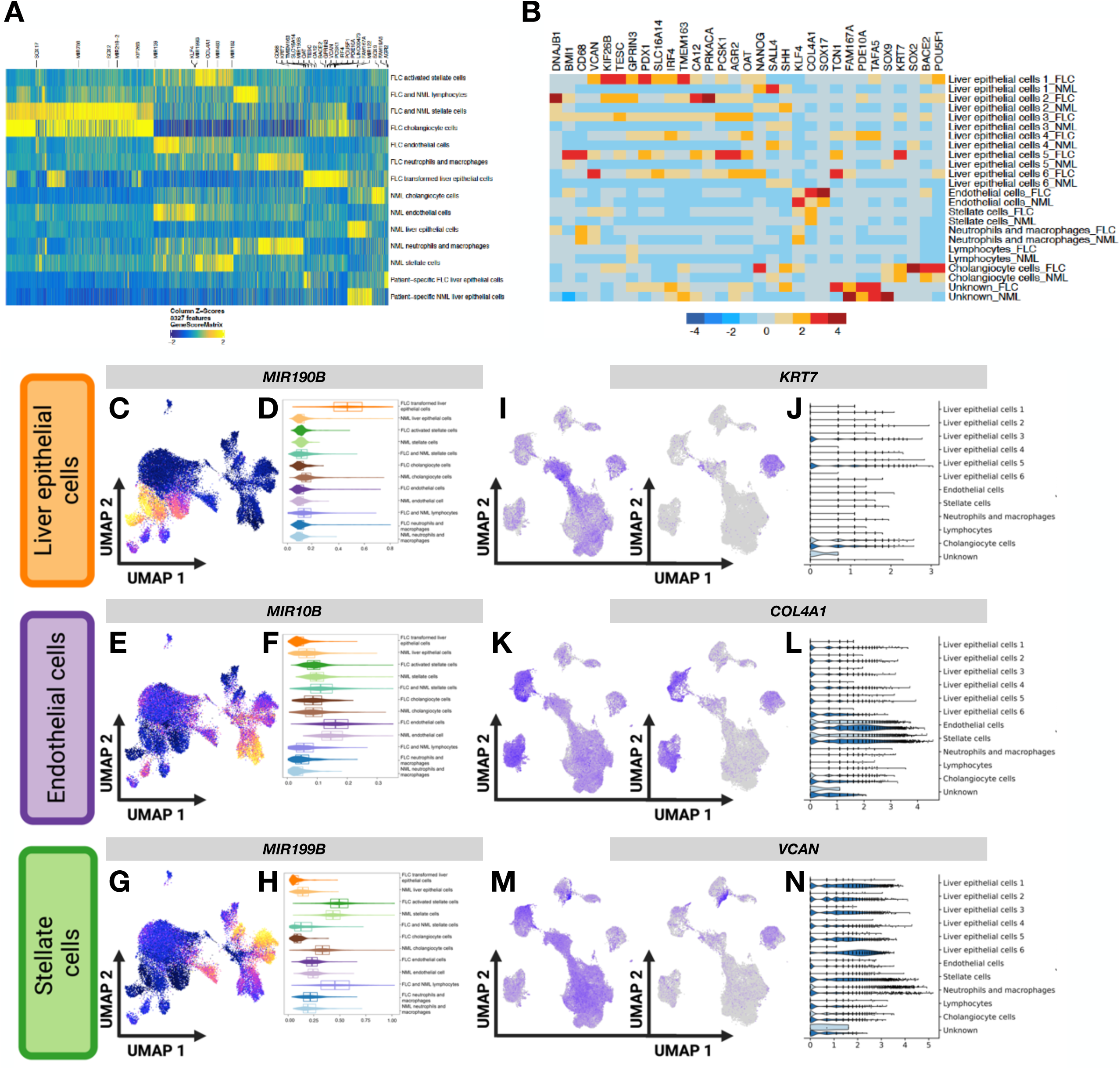
Single-nuclei ATAC-seq resolves cell type-specific molecular signatures in fibrolamellar carcinoma. **(A)** Heatmap depicting gene activity z-scores of enriched genes in each cell type in FLC and NML. Genes are labeled at the top of the heatmap. **(C, E, G)** ArchR gene activity score from snATAC-seq for three microRNAs upregulated in FLC (miR-190b, miR-199b, and miR-10b) showing different patterns of activity across cell types. **(D, F, H)** Violin plots depicting ArchR gene activity scores of selected FLC marker miRNAs. **(B)** Heatmap depicting z-score of average expression level for selected genes across each cell type in FLC and NML. **(I, K, M)** Signal from snRNA-seq for three genes of interest in FLC (KRT7, COL4A1, and VCAN) showing different patterns of expression across cell types. **(J, L, N)** Violin plots depicting gene expression of selected FLC marker genes. UMAP, Uniform Manifold Approximation and Projection; FLC, fibrolamellar carcinoma; NML, non-malignant liver.

Our previous bulk ChRO-seq work identified several transcription factor (TF) networks (*22*), including those of CREB1 and AP-1, that are likely completely rewired in FLC. However, the cell types driving that signal remain unknown. To address this knowledge gap, we examined the accessible chromatin landscape defined by our snATAC-seq data in FLC vs. NML tissue.

Specifically, we used MACS2 to perform peak calling followed by ArchR’s iterative overlap peak merging approach to generate a peak-by-cell matrix. This unveiled distinct marker peaks in each cell type from FLC and NML tissue (Fig. 3A), providing clearer stratification of cell types and tissue status than ArchR gene activity scores (Fig. 2A). Motif enrichment analysis within marker peaks using the cisBP database showed TF regulatory networks enriched in specific FLC or NML cell types (Fig. 3B). Notably, we detected significant enrichment of motifs for AP-1 TFs including JUN and FOS in marker peaks of not only FLC tumor epithelial cells but also FLC endothelial cells and FLC activated stellate cells relative to the NML counterparts (Fig. 3B). We also found marked loss of RXR-A/B and LXR-A/B (NR1H2/3) signal in FLC tumor epithelial cells compared to NML epithelial cells (Fig. 3B), which is consistent with loss of normal hepatocyte function. Detailed analysis of individual TFs showed that a subset of FLC endothelial cells drives the enrichment signal for ETV2, whereas a subset of FLC tumor epithelial cells are responsible for the CREB1 enrichment signal (Fig. 3C).

**Fig. 3.**
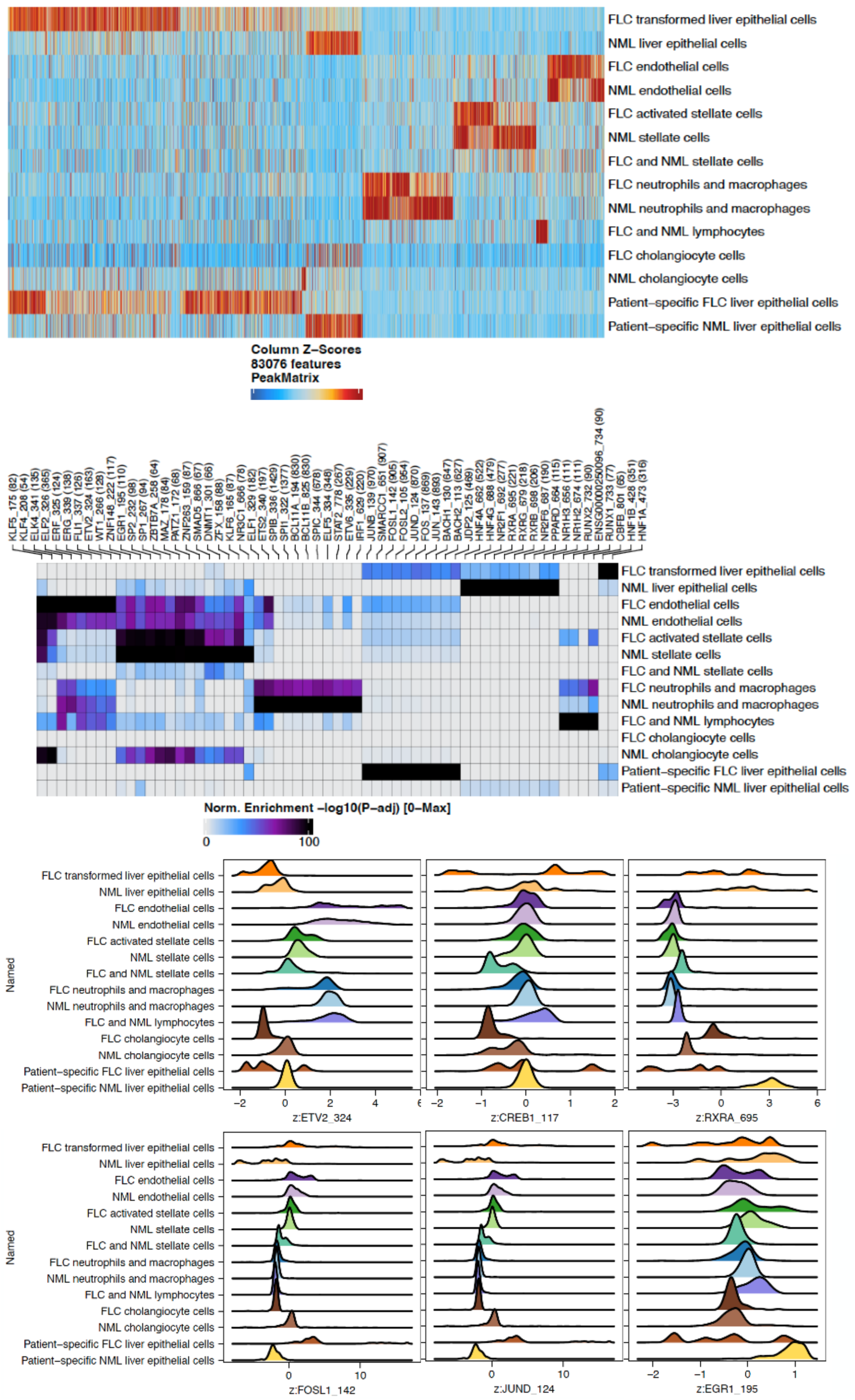
Single-nuclei ATAC-seq reveals cell-type specific transcription factor networks. **(A)** Heatmap depicting significantly enriched open chromatin peaks (as defined by chromatin accessibility) in each cell type in FLC and NML. **(B)** Heatmap depicting the top five transcription factor networks (based on motifs from cisBP) enriched in each cell type in FLC and NML. Transcription factors are labeled on the top. **(C)** Ridge plots depicting z-score density of motif presence for ETV2, CREB1, RXRA, FOSL1, JUND, and EGR1 across all cells in each cell type cluster (computed by ChromVAR). FLC, fibrolamellar carcinoma; NML, non-malignant liver.

### Inference of single-cell active enhancer landscape by deconvolution of bulk ChRO-seq data

In previous work we had defined the super enhancer landscape from bulk tissue. To infer the single-cell active enhancer landscape we conducted multi-omic data integrative analysis. This approach involves deconvoluting the bulk active transcription signal provided by ChRO-seq using single-cell accessible chromatin signal provided by snATAC-seq to determine the cell type profile of all distal enhancers. Using this approach, we determined the fraction of constituent enhancers within super enhancers that are present within the same cell type (Fig. 4A). Most previously defined super enhancers were fully or mostly intact in FLC tumor liver epithelial cells. There were also indications of ChRO-seq defined super enhancers that are partially but not fully intact in other cell types. For example, the gene versican (*VCAN*), which we previously reported as highly elevated in FLC (*33*) and functions in drug resistance (*34*), is associated with two super enhancers, one that is intact in FLC tumor liver epithelial cells and another that is present in both FLC tumor liver epithelial cells and FLC activated stellate cells (Fig. 4B). Upon analysis of the snRNA-seq dataset, we observed upregulated *VCAN* expression in both cell types (Fig. 4C). We then performed a more detailed analysis of *SLC16A14*, which we have shown previously is a unique marker of FLC. Both the *SLC16A14*-associated super enhancer (snATAC-seq) and mRNA expression (snRNA-seq) were strongly enriched (even nearly completely restricted) to FLC tumor liver epithelial cells (Fig. 4D and E), which has not been demonstrated previously.

**Fig. 4.**
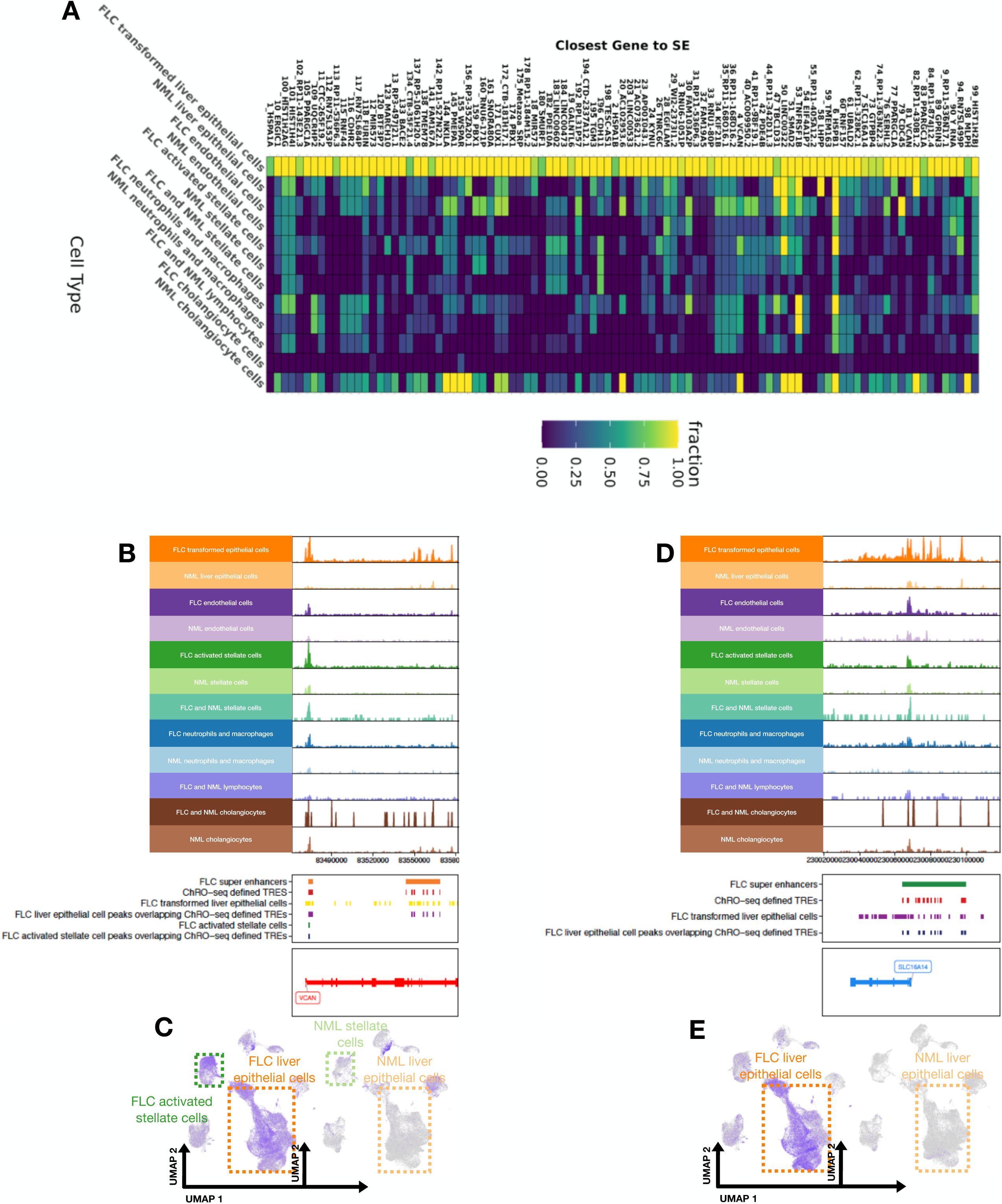
Integration of single-nuclei ATAC-seq, single-nuclei RNA-seq, and bulk ChRO-seq reveals cell type specific enrichment of super enhancers and their associated genes. **(A)** Heatmap depicting the fraction of overlap between single-nuclei ATAC-seq accessible regions called by MACS2 (q-value cutoff: 0.02) and regulatory elements within FLC-specific super enhancers (log2FC: 0.0, FDR: 0.01) called by dREG/ChRO-seq. **(B and D)** Browser track of local accessibility around (B) *VCAN* and (D) *SLC16A14* loci across all cell types in FLC and NML. **(C and E)** Single-nuclei RNA-seq UMAP overlay depicting expression of (C) *VCAN* and (E) *SLC16A14* across all cell types in FLC and NML. FLC, fibrolamellar carcinoma; NML, non-malignant liver.

### SLC16A14 exhibits dependency on fusion chimera DNAJB1-PRKACA protein

The marked specificity of SLC16A14 to tumor epithelial cells in FLC led to the hypothesis that the expression of this gene is dependent upon the expression of the FLC onco-driver *DNAJB1- PRKACA*. To test this hypothesis, we first evaluated by bulk RNA-seq whether *SLC16A14* is detected in other liver cancers that do not express *DNAJB1-PRKACA*. We confirmed that SLC16A14 exhibits dominant expression in FLC samples relative to HCC and CCA (Fig. 5A-C). We then performed knockdown of *DNAJB1-PRKACA* in a previously established FLC tumor epithelial cell line (*11*, *23*, *35*) by siRNA targeting at the fusion junction (*11*) and demonstrated significantly reduced *SLC16A14* expression by qPCR (Fig. 5D). Finally, we measured *SLC16A14* via qPCR in six different FLC cell lines derived from FLC primary or metastatic tissue, two of which spontaneously lost expression of *DNAJB1-PRKACA* upon passaging. We observed near complete loss of *SLC16A14* exclusively in the two cell lines that no longer expressed *DNAJB1- PRKACA* (Fig. 5E). Taken together, these findings provide evidence for the first time that robust *SLC16A14* expression in FLC tumor epithelial cells is dependent upon the activity of *DNAJB1- PRKACA*.

**Fig. 5.**
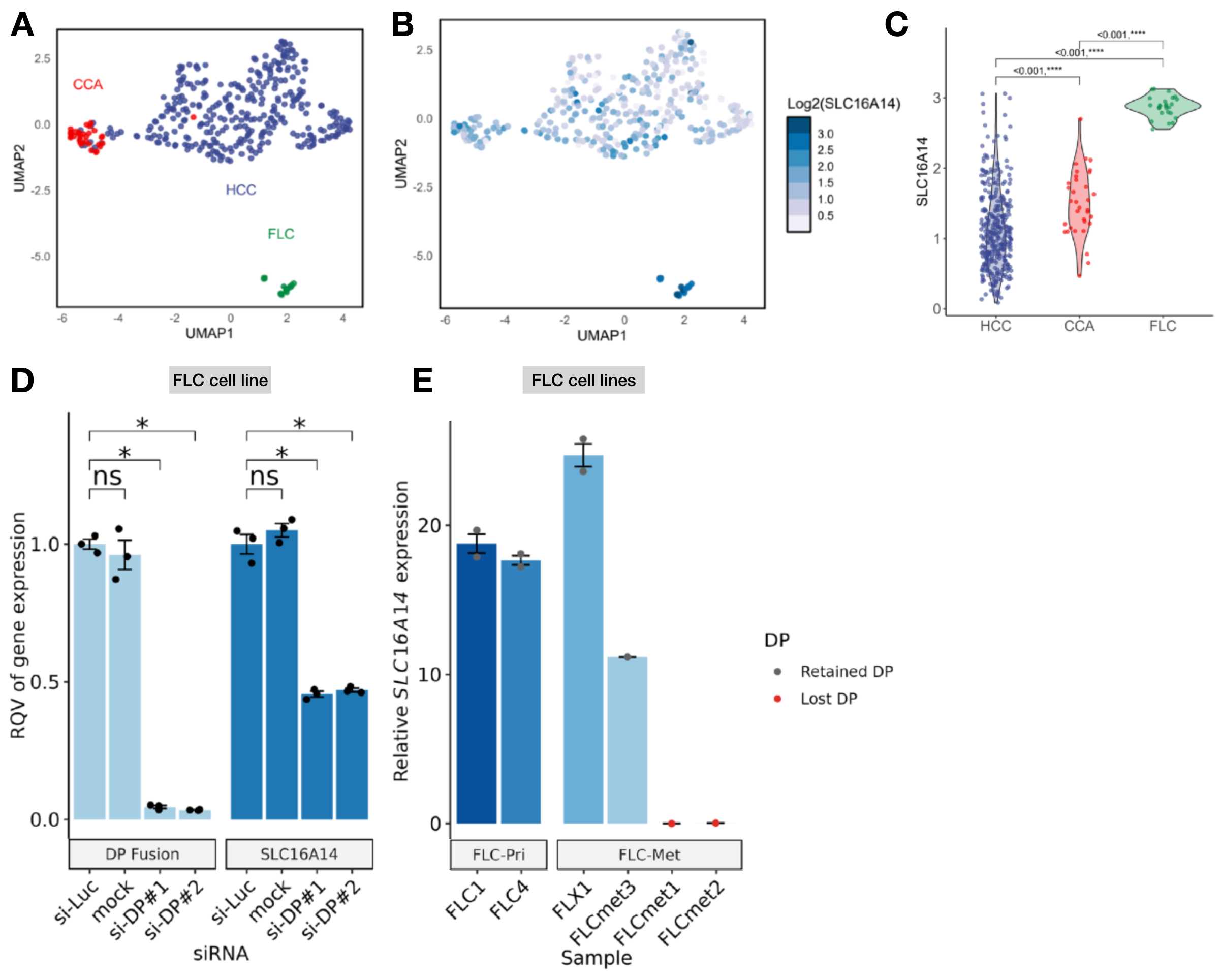
*SLC16A14* exhibits dependency on DNAJB1-PRKACA fusion expression. **(A)** UMAP visualization of gene expression profiles across four different types of liver cancers (FLC, green, n=20; HCC, blue, n=344; CCA, red, n=34. **(B)** *SLC16A14* expression by RNA-seq across all liver cancer types shown in panel A. **(C)** Violin plots of log2 expression of *SLC16A14* in HCC (blue), CCA (red), and FLC (green). **(D)** Relative gene expression of *DNAJB1-PRKACA (DP)* and *SLC16A14* assessed by qPCR in FLC cells (FLX1, n=3 independent experiments) after transfecting si-Luciferase (control), mock (no transfection), si-DP#1, or si-DP#2 (5nM). All data are normalized to *RPS9* and shown as mean ± SEM. Significance calculated with a non-parametric Wilcoxon signed-rank test. **(E)** Relative *SLC16A14* expression measured by qPCR in FLC primary cells (FLC1, FLC4, FLX1) or FLC metastatic cells (FLCmet3, FLCmet1, FLCmet2) and shown as mean ± SEM. n = 1 or 2 independent experiments for each biological replicate. FLCmet1 and FLCmet2 spontaneously lost DP expression. FLC, fibrolamellar carcinoma; HCC, hepatocellular carcinoma; CCA, cholangiocarcinoma.

### Cell and tissue models of FLC exhibit partial survival dependency on SLC16A14

The effect of SLC16A14 loss has only been examined in a cell model that was originally derived from the liver of transgenic mice overexpressing transforming growth factor alpha (*22*).

Therefore, we sought to test the role of SLC16A14 as a potential downstream effector of DNAJ-PKAc in human disease models of FLC via loss-of-function studies. Given our finding that SLC16A14 expression is essentially constrained to the epithelial compartment of FLC tumors, we first administered siRNA against *SLC16A14* in two distinct HepG2-DP+ clones (Fig. 6A), human liver epithelial cancer cell lines harboring the DNAJB1-PRKACA fusion by CRISPR/Cas9 deletion, mimicking the mutation in FLC patients. We observed significant reduction in *SLC16A14* expression levels in both clones. We then evaluated the effect of SLC16A14 knockdown on the viability of HepG2-DP+ cells. Our findings revealed a dose-dependent decrease in cell viability, with maximum efficacy (∼3-4 fold) observed at approximately 10nM concentration. We next sought to validate this finding in an FLC epithelial cell line (*11*, *23*, *35*) derived from primary tumor tissue. This experiment recapitulated both robust knockdown of *SLC16A14* (Fig. 6C) and concomitant and significant (∼2 fold) decrease in tumor cell viability (Fig. 6D). We further corroborated these results in a PDX-derived cell line (fig. S9). Finally, we conducted an experiment on live tissue slices from patient tissue. Upon administering siRNA against *SLC16A14* to tissue slices from NML, hepatocellular carcinoma (HCC), colorectal liver metastases (CRLM), and FLC, we observed a significant decrease in viability exclusively in FLC (Fig. 6E).

**Fig. 6.**
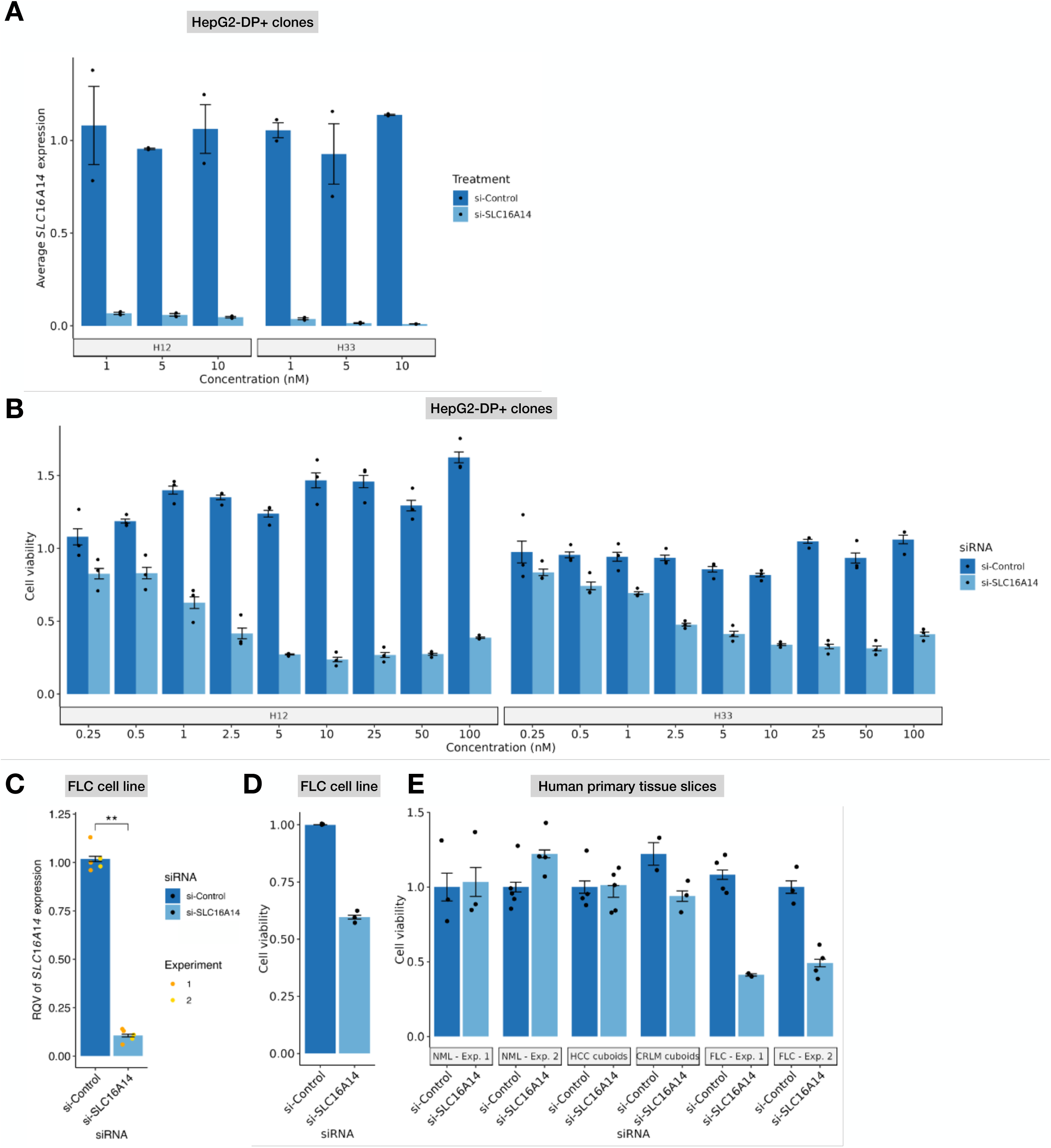
FLC cells exhibit unique dependency on SLC16A14. **(A)** Relative *SLC16A14* expression in HepG2-DP+ cell lines (H12 and H33, n=2 technical replicates for each clone at each concentration) after treatment with si-SLC16A14 (1nM, 5nM, and 10nM). All data relative to *GAPDH* and normalized to diH_2_O. **(B)** Cell viability in H12 (left) or H33 (right) HepG2-DP+ cells (n=3 technical replicates) after treatment with increasing amounts of si-SLC16A14 from 0.25nM to 100nM. **(C)** Relative *SLC16A14* expression in FLC cells (FLX1, n=3 technical replicates) after si-SLC16A14 treatment (30 pmol). All data relative to *RPS9*. Significance tested with non-parametric Wilcoxon signed-rank test. **(D)** Cell viability of FLC cells (FLX1, n=2 biological replicates) after si-SLC16A14 treatment (30 pmol). **(E)** Cell viability of cuboids from tissue slices of primary human NML, HCC, CRLM, or FLC tissue after treatment with si-SLC16A14 (30 pmol). Each bar plot represents a biological replicate, and data points represent technical replicates. n=3-5 technical replicates for each biological replicate. FLC, fibrolamellar carcinoma; NML, non-malignant liver; HCC, hepatocellular carcinoma; CRLM, colorectal liver metastatic tissue.

### Inference of novel cell-cell communication in FLC

To explore potential intercellular communication mechanisms in FLC, we analyzed the snRNA-seq dataset to define newly activated (Fig. 7) and rewired (Fig. 8) ligand-receptor pathways using CellChat (*36*). This tool was chosen for its intuitive interface and extensive database of signaling pathways. First, we found that the number and strength of ligand-receptor interactions are dramatically altered in FLC relative to NML (Fig. 7A), most notably the involvement of endothelial cells, stellate cells, and cholangiocytes. Communication among epithelial cell subtypes is also rewired in FLC (Fig. 7A). We next examined the information flow of each signaling pathway in FLC and NML, identifying several NML- and FLC-specific pathways (Fig. 7B). Notably, six FLC-specific pathways emerged: FGF, BRADYKININ, PERIOSTIN, GDF, CALCR, and EGF. Further analyses revealed very different patterns of relevant cell types for these pathways of interest (Fig. 7C, 7D). We then selected three of the FLC-specific pathways— GDF, CALCR, and FGF—for more detailed investigation (Fig. 7E-J and fig. S8). The GDF signaling network revealed a single ligand-receptor pair, GDF15- TGFβR2 (fig. S10A).

**Fig. 7.**
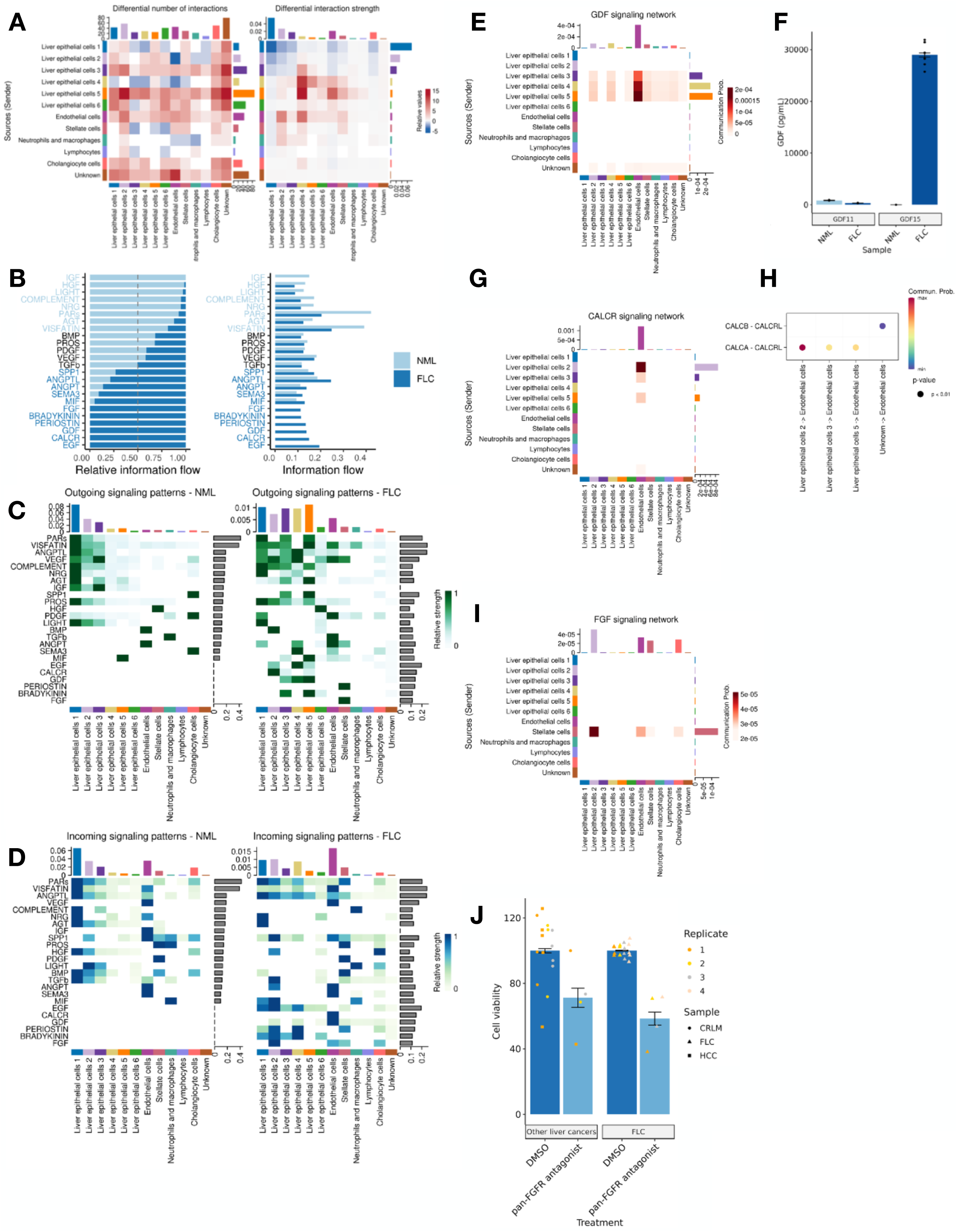
CellChat reveals novel cell-to-cell communication pathways in FLC. **(A)** Heatmap depicting the differential number of interactions (left) or interaction strength (right) between FLC and NML. Red corresponds to more in FLC, blue corresponds to more in NML. Adjacent to each heatmap, the right bar plot depicts the sum of row values for outgoing signals and the top bar plot depicts the sum of column values for incoming signals. **(B)** Information flow showing the sum of communication probability (total weights) for shared or NML-or FLC-specific pathways. Signaling pathways enriched in NML are shown in light blue, and signaling pathways enriched in FLC are shown in dark blue. **(C and D)** Outgoing **(C)** and incoming **(D)** signaling patterns for NML (left) and FLC (right) of pathways identified in both datasets. Adjacent to each heatmap, the bar plots on the right depicts the sum of row values for communication probability for that pathway. The bar plots on top shows the sum of column values for signaling strength for specific cell types. Pathways are labeled on the left, and cell clusters are labeled at the bottom. **(E, G, and I)** Heatmap depicting GDF **(E)**, CALCR **(G)**, and FGF **(I)** signaling pathways in FLC. Source (sender) cells are labeled on the left, and the receiver cell type(s) is/are labeled on the bottom. The right bar graph depicts sum of contribution from sources. The bar graphs on top depict the sum of column values for receptors. **(F)** Secreted GDF11 and GDF15 levels in media for biological replicates of NML cuboids from patient tissue slices (GDF11, n=2; GDF15, n=2) or FLC cells (GDF11, n=6; GDF15, n=6). **(H)** Bubble plot depicting significant ligand-receptor pairs within the CALCR pathway. Ligand-receptor pairs are labeled on the left, and source-receiver cell types are labeled on the bottom. **(J)** Cell-titer glo measuring cell viability in various liver cancers or FLC after administering pan-FGFR antagonist (2.5nM). Replicates shown in different colors, cancer types shown in different shapes. FLC, fibrolamellar carcinoma; NML, non-malignant liver; HCC, hepatocellular carcinoma; CRLM, colorectal liver metastatic tissue.

**Fig. 8.**
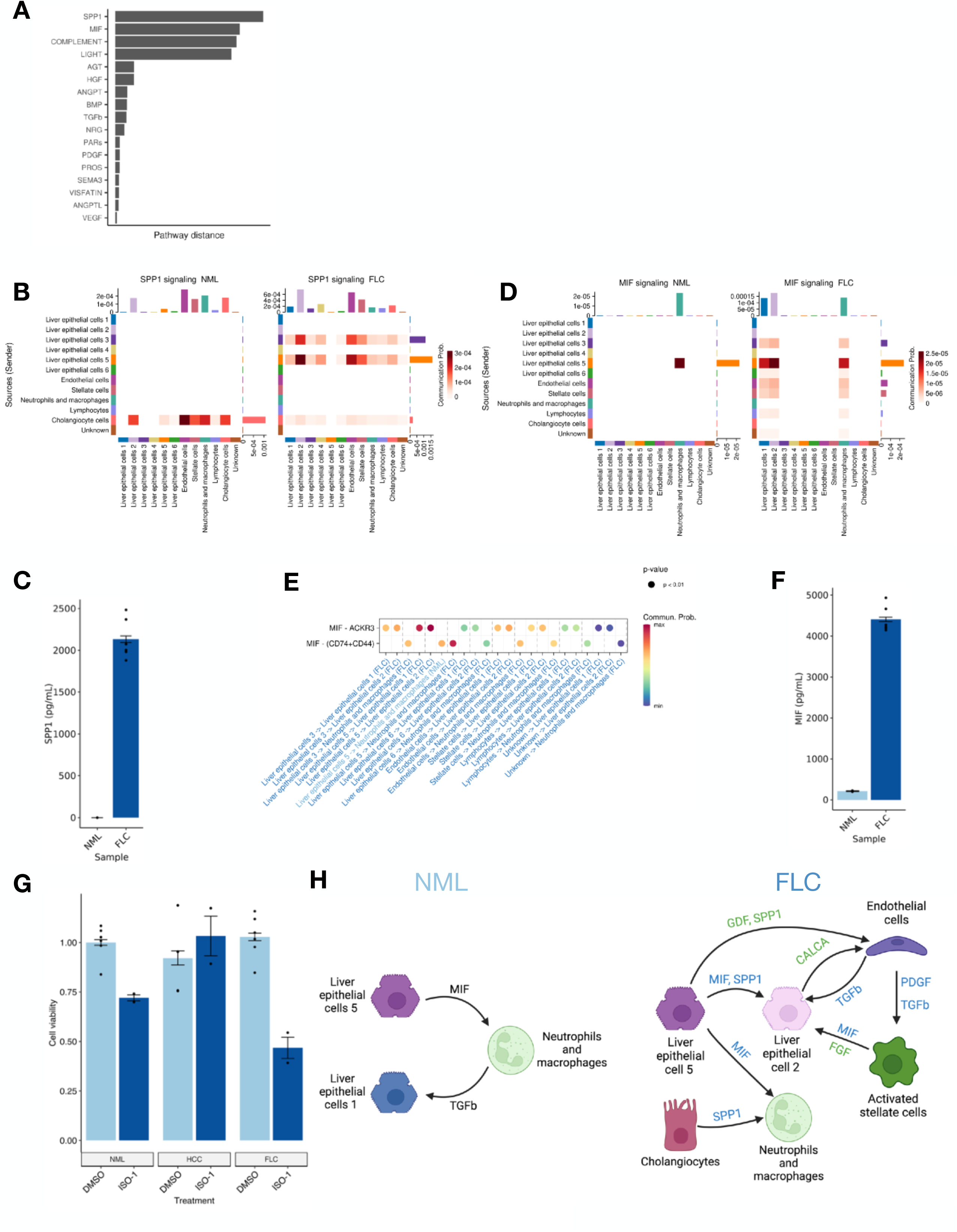
CellChat reveals cell-to-cell communication patterns that are dramatically rewired in FLC compared to NML. **(A)** Extent of rewiring (pathway distance) in FLC of signaling pathways that are shared between FLC and NML. **(B and D)** Heatmap depicting rewiring of SPP1 **(B)** and MIF **(D)** in NML (left) compared to FLC (right). Source (sender) cell types are labeled on the left, and the receiver cell types are labeled on the bottom. The right bar graph depicts sum of contribution from sources. The bar graphs on top depict the sum of column values for receptors. Intensity of heatmap reflects communication probability. **(C and F)** Secreted SPP1 **(C)** and MIF **(F)** levels in media for biological replicates of NML cuboids from patient tissue slices (n=2), or FLC cells (n=6). **(E)** Comparison of the significant ligand-receptor pairs in the MIF pathway in NML (light blue) or FLC (dark blue). Rows depict specific ligand-receptor pairs and columns define cellular sources and receptors. Dot color reflects communication probabilities and dot size represents computed p-values. **(G)** Cell viability of cuboids from tissue slices of primary human NML, HCC, or FLC tissue after treatment with ISO-1 (1 uM). Each bar plot represents a biological replicate, and data points represent technical replicates. n=2-6 technical replicates for each biological replicate. **(H)** Schematic summarizing rewiring and activation of the most highly significant ligand-receptor pairs in FLC (right) compared to NML (left). Pathways (ligands) labeled in green are FLC-specific pathways; pathways (ligands) labeled in blue are shared between FLC and NML but dramatically rewired in FLC.

Specifically, GDF15 is sourced predominantly from liver epithelial cells 3, 4, and 5, and its receptor TGFβR2 is mainly expressed within endothelial cells (Fig. 7E). This finding leads to the hypothesis that GDF15 is a highly secreted molecule from FLC tumor cells. To test this hypothesis, we measured secreted proteins in NML tissue and FLC cell cultures and observed a significant increase in GDF15, but not GDF11, secretion in FLC compared to NML, consistent with CellChat results (Fig. 7F) as well as bulk RNA-seq data (fig. S10B and C).

The CALCR pathway exhibited the second-highest signaling strength in FLC (Fig. 7B), with the strongest connection between liver epithelial cell 2 and endothelial cells (Fig. 7G, H). The liver epithelial 2 compartment serves as the primary source of the CALCA ligand (fig. S10D) and we found that CALCRL is the most dominant receptor in FLC, expressed almost exclusively in endothelial cells (fig. S10D). We note that CALCA is upregulated in FLC as measured bulk RNA-seq also (fig. S10E), as well as specific to FLC compared to other common liver cancers according to TCGA data analysis (fig. S10F).

Lastly, for the FGF pathway, which is shown to be upregulated and specific to FLC (fig. S10G-I), our analysis inferred strong communication between activated stellate cells and liver epithelial cell 2 (Fig. 7I), the same epithelial cell subtype that expresses CALCA. Given the predicted signaling in the epithelial compartment, we hypothesized that the FGF pathway may directly control tumor cell viability. We assessed the functional impact of a pan-FGFR antagonist in tissue slices from several different liver cancers and found that there was a significant effect on FLC, albeit only slightly more so than on other liver cancer types (Fig. 7J).

### Rewired cell-cell communication in FLC relative to NML

We then focused on the rewired (as opposed to newly activated) secretory pathways in FLC relative to NML. We identified several pathways with significant rewiring (or “pathway distance” between FLC and NML), such as SPP1 and MIF, which have not previously been reported in FLC studies, or TGFβ, which has been highlighted before but not at single-cell resolution (Fig. 8A).

For the SPP1 pathway, five ligand-receptor pairs remain active, and we infer the deactivation of the SPP1-(ITGAV+ITGB3) and the activation of SPP1-(ITGAV-ITGB6) in FLC relative to NML (fig. S11A). Examination of our previously published bulk RNA-seq data confirms that SPP1 is elevated in FLC relative to NML, and in FLC relative to HCC, but lower than CCA (fig. S11C and D). The dramatic rewiring of SPP1 signaling arises from the shift in source-receptor cell types. In NML, SPP1 appears primarily sourced from cholangiocytes, with receptors expressed in various cell types, most notably endothelial cells (Fig. 8B). However, in FLC, the primary source of SPP1 is tumor epithelial cells, particularly liver epithelial 5, with receptors expressed most dominantly in liver epithelial 2. We also noted the dramatic shift in the SPP1-CD44 interaction, observing that the major source changes from cholangiocytes in NML to tumor epithelial cells in FLC (fig. S11E and F). These findings motivate the hypothesis that SPP1 is a highly secreted molecule in FLC tumors. To test this hypothesis, we assessed SPP1 among the secreted proteins in NML tissue and FLC cell culture and observed a significant increase in SPP1 secretion in FLC (Fig. 8C).

We also examined the MIF pathway, which in NML is limited to liver epithelial cell 5 as the source, with NML neutrophils and macrophages as the sole receivers, mediating the MIF- (CD74+CD44) ligand-receptor pair signaling (Fig. 8D). In contrast, in FLC we observed an increase in the number of sources, primarily FLC liver epithelial 3, 5; FLC endothelial; and FLC stellate cells, diversifying the MIF-(CD74+CD44) interaction (Fig. 8E). Further inspection reveals unique activation of the MIF-ACKR3 ligand-receptor pair in FLC (fig. S11G and H).

Examination of our previously published bulk RNA-seq data revealed that ACKR3 (CXCR7) is significantly elevated and unique in FLC compared to HCC and CCA (fig. S11I and J). Another noteworthy observation is that ACKR3 expression shifts from immune cells in NML to liver epithelial cells (subtype 2) in FLC (Fig. 8E). Like with SPP1, MIF is significantly increased as a secreted factor from FLC cells compared to NML (Fig. 8F). We further interrogated MIF signaling by administering MIF inhibitor ISO-1 to live tissue slices from patient tumors (Fig. 8G). We observed a significant reduction in cell viability in FLC but not change in HCC.

As for TGFβ signaling, both ligand-receptor pairs present in NML, TGFβ1-(TGFβR1+TGFβR2) and TGFβ1-(ACVR1B+TGFβR2), undergo dramatic changes in FLC (fig. S12A, B, and C).

Specifically, we observed that neutrophils and macrophages as the source in NML shifts to endothelial cells as the source in FLC, with multiple different cell types receiving the signal (fig. S12D and E). A third ligand-receptor pair TGFβ1-(ACVR1+TGFβR2) is activated and is also sourced from FLC endothelial cells (fig. S12C, F, and G). Several of the main findings from the CellChat analysis are summarized in Fig. 8H, revealing completely new and rewired pathways that highlight candidate druggable targets in FLC.

## Discussion

FLC remains a deadly rare form of liver cancer with limited therapeutic options, underscoring the urgent need to decode its tumor biology for the identification of potential therapeutic targets. Through single-cell sequencing technologies applied to primary FLC tumors and adjacent non-malignant liver (NML) samples, we aimed here to deconvolute previously reported signals, offering novel insights into FLC tumor biology that were previously obscured by bulk tissue omic approaches. Our study represents the first single-cell, multiomic investigation of FLC to date. We anticipate that this dataset will serve as a valuable resource, aiding in deeper comprehension of FLC tumor biology and inspiring the development and testing of new treatment strategies. Furthermore, we envision this study to serve as a reference for scientists in the field seeking to understand FLC intra-tumoral heterogeneity, as well as to drive hypotheses about new therapeutic avenues that merit further investigation.

Our study deepens and broadens our understanding of FLC tumor biology. We have previously demonstrated that distal transcriptional regulatory elements (TREs) stratify FLC from NML more effectively than gene expression alone (*22*), consistent with findings indicating that enhancer activity provides greater sensitivity in delineating sample status (*37–40*). Our analysis further reveals that the dimensionality reduction strategy applied to snATAC-seq by ArchR identifies cell types delineated by tissue status more effectively than the PCA approach applied to snRNA-seq by Seurat. This observation likely stems from ArchR’s unique iterative-LSI dimensionality reduction approach, which considers accessible features acting as distal TREs.

Both single-cell approaches unveil the cellular composition of FLC for the first time. SnATAC-seq and snRNA-seq both highlight a substantial increase in the relative abundance of stellate cell and endothelial cells in FLC vs. NML, likely reflecting increased proliferative capacity of both populations in the tumor setting. The fibrotic bands characteristic of FLC may be partially attributed to several factors that we identified as specific to FLC activated stellate cells, including CTGF/CCN2, a well-established pro-fibrotic factor, and miR-199b, family members of which have been shown to activate the TGFβ pathway (*41*). Also, we showed that miR-10b, one of the most highly elevated miRNAs in FLC (*23*), is altered not in the tumor epithelial cells as previously thought, but rather primarily in endothelial cells in the tumor microenvironment.

The transcription factor (TF) network analysis reveals increased activity in FLC endothelial cells of ETV2, a TF known to mediate endothelial transdifferentiation (*42*) (and thereby promoting the development of a metastatic niche) in glioblastoma, and likely also contributing to the invasive nature of FLC. Additionally, TF network analysis shows that the rewired CREB1 transcriptional network in FLC is driven largely by the tumor epithelial compartment, consistent with CREB1 being downstream of PKA signaling (*43*, *44*), which is dysregulated in FLC due to DNAJ-PKAc activity in tumor epithelial cells.

Our study also presents a novel integrative approach to infer cell-type specific super enhancers (SEs). By leveraging chromatin accessibility patterns in single cells, we successfully deconvolute bulk chromatin activity previously characterized using ChRO-seq using ChIPpeakAnno (*45*).

Incorporating expression data from snRNA-seq, we demonstrate that high expression of several FLC markers occurs only in those cell types in which the associated SEs are largely intact. We show that the most robust FLC markers to date, *SLC16A14* and its associated SE, are essentially specific to FLC tumor epithelial cells. Moreover, we provide the first evidence to date in cell models of FLC that *SLC16A14* expression is dependent on *DNAJB1-PRKACA* expression. To assess the potential dependence of FLC on SLC16A14, we conduct a series of *SLC16A14* knockdown (siRNA) experiments in three different models of disease: HepG2-DP+ clones, an FLC epithelial cell line, and FLC primary tumor tissue slices. Our investigations reveal that knockdown of *SLC16A14* consistently leads to significantly decreased cell viability across all three models. Taken together these results confirm SLC16A14 as a downstream mediator of DP and represent the first functional assessment of FLC dependency on SLC16A14 that warrants further investigation in future studies.

With the snRNA-seq dataset we delve deeper into novel and rewired cell communication pathways in FLC relative to NML using CellChat. Notably, we highlight the potential involvement in FLC of GDF15 signaling, shown to drive cell growth, migration, and invasion in some other cancers (*46*), as well as also inhibit T cell migration in melanoma. Additionally, we observe robust activation of FGF signaling network in FLC. In other cancers FGF signaling partners with RAS/MAPK, PI3/AKT, and/or PLCγ pathways (*47*). Given previous observations of aberrations in the RAS/MAPK pathway in FLC, we hypothesize that these connections are also relevant in FLC. Consistent with this notion, treatment of FLC primary tumor tissue slices with pan-FGFR antagonists significantly reduce viability. Finally, we identify significant rewiring in the SPP1, MIF and TGFβ signaling pathways in FLC relative to NML. The SPP1 and MIF pathways diversify their targets in an FLC background, and both converge on macrophages and neutrophils. Both of these pathways have been shown to modulate the tumor microenvironment to suppress the immune response (*48*, *49*). Furthermore, the MIF-ACKR3 (CXCR7) (*48*) interaction appears relevant in tumor epithelial cells in FLC whereas it is likely active only in immune cells in NML. We propose that MIF signaling through CXCR7 promotes FLC tumor cell proliferation and migration. Regarding TGFβ rewiring, we observe a shift from neutrophils and macrophages as a source in NML to endothelial cells in FLC. Because endothelial cells are much greater in abundance relative to neutrophils and macrophages in FLC, this finding points to a likely large increase in the strength of TGFβ signaling overall in FLC. Collectively, these findings offer new insights into signaling in FLC tumors and highlight potential new therapeutic targets to test in the future.

Despite the substantial advances to our understanding of FLC tumor biology brought about by these single-cell multi-omic analyses, there are several limitations that warrant attention. Firstly, only some of the samples in our study were patient-matched; in the future it will be important to increase the number of matched samples to adequately account for patient-specific confounding factors. Secondly, the use of separate assays to profile accessible chromatin and gene expression introduces potential confounders. A true multiomic platform profiling both accessibility and gene expression within a single cell would address this limitation. Finally, our study constitutes a “pseudo” multiomic integration between snATAC-seq and snRNA-seq; a true multiomic integration would allow for a stronger association between distal peak accessibility and TF networks determined by snATAC-seq with gene expression determined by snRNA-seq.

Nonetheless, we believe that this study contributes substantially to our understanding of FLC biology and provides a very valuable resource for the field studying this rare cancer and related malignancies.

## Materials and methods

### snATAC library preparation

#### Tn5 storage and transposome assembly

The Tn5 transposase was prepared using a modified protocol derived from Hennig B.P.’s method (*50*) and stored at-80°C in a storage buffer containing 100 mM HEPES-KOH at pH 7.2, 0.2 M NaCl, 0.2 mM EDTA, 2 mM DTT, 0.2% Triton X-100, and 20% glycerol. The Tn5 transposase (5 µM) was diluted by adding one volume of 80% glycerol. Subsequently, transposomes were assembled by adding 0.1 volume of Tn5 adaptors (25 µM) to the diluted Tn5 (2.5 µM). The mixture was incubated at room temperature for 12–24 hours. The resulting transposomes (∼2 µM) can be used directly or stored at-20°C.

#### Tagmentation

The nuclei isolation and tagmentation were conducted following a previously described protocol by Corces, M (*51*). The modification in the isolation steps includes two homogenization stages: initially in 1x Homogenization Buffer (HB) containing 5 mg/ml Collagenase Type IV using a loose pestle, followed by the second homogenization step in 1x Washing buffer (1x ATAC-RSB + 0.1% Tween-20). Combinatorial single-nucleus barcodes were generated using a strategy modified from and developed in the Genomics Innovation Hub at Cornell. Nuclei suspension (8 μL) was distributed onto 96-well plates, and 1 μL of each i5 and i7 transposome was added to each well, resulting in 96 combinations of Tn5 barcodes per plate. Each well contains ∼5000 nuclei and 400 µM transposomes in 1X TD buffer (10 mM Tris-HCl, pH 7.6, 5 mM MgCl2, 10% DMF, 0.33X PBS, 0.1% Tween20, 0.01% Digitonin). The tagmentation reaction plate was incubated at 50°C for 30 minutes, and the reaction was terminated by adding 10 μL 20 mM EDTA (15 minutes at 37°C). Next, 20 μL of sorting buffer (1× SB: 1× PBS, 2 mM EDTA, 20 ng/mL BSA) was added to each well, and nuclei were repooled into a single 5 ml tube. Nuclei were then stained with DRAQ7 for 15 minutes (ABCam, ab109202), passed through a 30 μm filter, and reisolated by FACS using a FACSMelody instrument (Becton, Dickinson). A 96-well destination PCR plate was preloaded with 10 μL of modified sorting elution buffer (1× SEB: 10 mmol/L TRIS pH 8.0, 12 ng/μL BSA, 0.05% SDS), 25 nuclei were distributed into each well, and incubated for 10 minutes at 55°C to disrupt Tn5 and finish tagmentation. We added 2.5 μL of 5% Triton-X100 per well to neutralize the SDS prior to PCR.

#### PCR amplification and snATAC library preparation

For each well in a 96-well plate, the 12.5 µL of 2X PCR reaction master mix contains 5 µL Q5 Reaction buffer, 5 µL High GC Enhancer, 0.5 µL dNTP mix (10 mM), 0.25 µL Q5 DNA Polymerase, 0.5 µL Universal i5 primer (25 µM), 0.75 µL Nuclease-Free Water, and a unique barcoded i7 primer(final concentration 500nM) were added. Libraries were amplified using the following program: 72°C for 5 min, 98°C for 30 sec, then 15 cycles of (98°C for 15 sec, 66°C for 30 sec, 72°C for 40 sec). PCR products from all wells were combined and purified using QIAquick spin columns, followed by two rounds of size selection with AMPure XP beads(Beckman Coulter, A63880). The first round is at 0.5 X/1.2 X, and the second round is at 1.2 X. Finally, the DNA was eluted in 40 µL of 10 mM Tris, pH 8.0.

#### Quantification and sequencing of snATAC libraries

The concentration of library was measured using the Qubit. snATAC-seq libraries are between 3 ng/μL and 10 ng/μL. DNA fragment length distribution was analyzed using the Bioanalyzer High Sensitivity DNA Chip (Agilent Genomics). Libraries were sequenced on the Illumina HiSeq X platform (3 batches; 4 lanes total) at Novogene.

### ArchR pipeline for single-nucleus ATAC (snATAC) analysis

#### snATAC-seq preprocessing

Demultiplexed fastq files were processed using cutadapt, UMI tools, and custom scripts to parse and assemble combinatorial barcodes from read segments and extract valid single-nucleus barcodes (with error correction). Preprocessed reads were mapped to the human genome (hg38) with bwa-mem and duplicates removed with UMI tools. Deduplicated bam files were sorted and indexed using samtools and converted to fragment files using the single-cell analysis tool kit Sinto (https://github.com/timoast/sinto) with –use_chrom “” and –barcode_regex “(?⇐_)(.*)(? =_)” parameters. Fragment files are then sorted and finally used to generate tabix files using tabix prior to loading into ArchR.

#### snATAC-seq QC and dimensionality reduction and clustering analysis

The transcription start site (TSS) enrichment score and fragment number of each nucleus is calculated using ArchR (*29*) v1.0.1. Nuclei with TSS enrichment score less than 3 and fragment number less than 1,000 are removed. Doublet scores were calculated with default parameters.

Briefly, for the snATAC-seq cohort, we employed ArchR to construct a matrix of contiguous fragments, quantifying insertions within. We then utilized ArchR’s iterative latent-semantic indexing (LSI) for dimensionality reduction on the top 25,000 most variable genomic tiles. In analyzing the snRNA-seq cohort, we utilized Seurat for principal component analysis (PCA)- based dimensionality reduction, focusing on the top 3,000 most variably expressed genes.

Subsequently, cells were categorized into transcriptionally distinct clusters using the first 30 principal components (PCs). We then used the default harmony algorithm to correct for differences caused by batch effects and added clusters using the “addClusters” function.

#### Identification of marker features

We identified cluster markers using the function “getMarkerFeatures” with default parameters and then applied the “addImputeWeights” function to impute the weights of markers. We visualized marker features using the ‘plotEmbedding’ function for UMAPs or ‘plotGroups’ for violin plots. Heatmaps depicting FLC markers were visualized using the ‘plotMarkerHeatmap’ function with log2FC >= 1, FDR <= 0.2 thresholds.

#### Peak calling and data visualization

To call peaks in snATAC-seq, pseudo-bulk replicates were generated using default parameters for ‘addGroupCoveragè for each inferred cell type and pseudo-bulk peak calling was conducted using MACS2 using the ‘addReproduciblePeakSet’ function and a custom argument cutOff = 0.2. Peak calls were then merged into a single profile using ArchR unique iterative overlap approach.

#### Differential peak accessibility and transcription factor network analysis

Enriched peaks in each cluster were determined using default parameters of the ‘getMarkerFeatures’ function, identifying the most enriched peaks in a given cluster relative to all other clusters. A marker peak heatmap was plotted using the ‘plotMarkerHeatmap’ function using log2FC >= 1, FDR <= 0.2 thresholds. Marker peaks were used to analyze transcription factor networks using a motif database supplied by cisBP. We used ChromVar to visualize transcription factor motif enrichment distributions on a per-cell-basis using the ‘addMotifsAnnotations’ function using log2FC >= 1, FDR <= 0.2 thresholds with optional parameter ‘motifSet = “cisbp”.

#### Peak overlap analysis

Super enhancer intactness was inferred by conducting an overlap analysis between accessible snATAC-seq peaks called by MACS2 (default parameters) and ChRO-seq defined TREs defined by dREG (log2FC >= 0, FDR <= 0.05), using the R package ‘ChIPpeakAnnò. Accessible and TRE peaks are denoted as overlapping if there is at least one base overlap. Super enhancer intactness is reported as the fraction of overlapping ChRO-seq and snATAC-seq peaks divided by the total number of constituent ChRO-seq defined TREs that make up the super enhancer. Heatmap was constructed using ggplot2’s geom_tile function.

#### Plotting browser tracks

To plot browser tracks, we used the “plotBrowserTrack” function and arranged track rows from highest to lowest accessibility using the “useGroups” parameters.

### SnapATAC2 pipeline for single-nucleus ATAC (snATAC) analysis

Box plots and stacked bar plots were created using seaborn (*52*) (v0.13.2) and matplotlib (*53*) (v3.8.2). Unless mentioned, functions were used with default parameters.

#### snATAC-seq QC and dimensionality reduction and clustering analysis

The bed files were imported after sorting into snapATAC2 (*31*) (v2.5.3) and cells with less than 1000 fragments were filtered out. The following QC was performed on a per sample basis. The transcriptional start site enrichment (tsse) score was calculated using snapATAC2 with the function ‘tsse’. Cells with less than 1000 counts and with a tsse score bellow 5 were filtered out. The tile matrix was then generated using a bin size of 500 and the top 250,000 most variable features were kept. Doublet scores were generated using 30 principal components with the function ‘scrublet’ and cells with a doublet probability over 0.2 were removed. The data were then combined and again the top 250,000 most variable features were selected for downstream analysis. Dimensionality reduction was performed using snapATAC2’s matrix-free spectral embedding algorithm, keeping the first 50 components. Batch correction was then performed using the ‘harmony’ function. The neighborhood graph was generated using the harmonized data and setting the function ‘knn’ to 50 nearest neighbors. To visualize the data, Leiden clustering was performed at different resolutions with the function ‘leiden’ and plotted on a UMAP with the ‘umap’ function. Clusters that were patient specific were subsetted out and the data was re-clustered.

#### Identification of marker features

Using the function ‘make_gene_matrix’ we generated a cell by gene activity matrix by counting the Tn5 insertions in gene body regions. Using Scanpy (*54*) (v1.9.8) genes expressed in less than 5 cells were filtered and the counts were normalized and logarithmized. To denoise the data and recover transcripts ‘magic’ was run through Scanpy. For each cluster, differentially expressed genes were determined using a Wilcoxon test with the function ‘rank_genes_groups’ and were subsequently used to determine cell types.

#### Peak calling, identification of marker regions and differentially accessible regions, and data visualization

To call peaks in the snATAC-seq data, pseudo-bulk replicates were created using ‘merge_peaks’ for each inferred cell type and pseudo-bulk peak calling was performed using ‘macs3’ in snapATAC2. Then a cell by peak matrix was generated using the ‘make_peak_matrix’ function. Marker regions for each cell type were identified using the function “marker_regions”. To find differentially accessible regions (DARs) between two cell types, the ‘diff_test’ function was used.

##### snRNA library preparation

#### Single nuclei isolation

Single nuclei were isolated from FLC tumors or matched surrounding liver tissue as previously described (*55*), with some modifications. Briefly, frozen tissue samples were briefly hand-thawed, washed in ice-cold PBS, and cut into small pieces using dissecting scissors. All subsequent steps were performed directly on wet ice. Tissue was homogenized to liberate nuclei in TST buffer(*55*) with 20 strokes of Dounce A and B homogenizers, and adequate nuclei isolation was confirmed by trypan blue uptake in a small aliquot of lysate. Lysate was 40μm filtered and recovered in ST buffer(*55*) containing 0.5% BSA. Nuclei were pelleted in a swinging rotor centrifuge at 500g for 5 minutes at 4°C. Supernatant was then decanted and nuclei gently resuspended in PBS (no Mg/Cl) containing 1% BSA and a 1:200 dilution of Protector RNase (Sigma Aldrich PN-3335399001) for counting. All subsequent processing and single-cell emulsion was per kit instructions, Chromium Next GEM Single Cell 3’ Reagent Kits v3.1 revision D (10X genomics) on a 10X Chromium Controller.

#### snRNA-seq library generation and sequencing

Sequencing libraries were generated per kit instructions, Chromium Next GEM Single Cell 3’ Reagent Kits v3.1 revision D (10X genomics), with the only modification being cDNA (step 2.2) was amplified 2 cycles more than recommended to account for the lower cDNA yield from nuclei. Library size and quality was determined on an Agilent Tapestation 2200. All samples passing QC were submitted for deep sequencing by GeneWiz with a targeted read depth of 20,000 reads per cell.

#### snRNA-seq preprocessing

Fastq files were generated from the single-nuclei sequencing raw data and aligned to the human genome (hg38) via CellRanger (v.7.0.1). Transcriptomic signal originating from productive gel beads-in-emulsion rather than ambient cell-free transcripts was ensured by processing raw and filtered gene/count matrices produced by CellRanger through SoupX default parameters of the autoEstCont method.

### Seurat pipeline for single-nucleus RNA (snRNA) analysis

#### snRNA-seq QC and dimensionality reduction and clustering analysis

Corrected matrices after ambient RNA removal were imported and initialized in Seurat (v5.0.1) for all downstream quality control and analysis steps. First, doublets were computationally predicted and removed by scDblFinder. Low quality nuclei with ˂1000 genes detected or ˃20% of reads mapping to mitochondrial genes were filtered out. The remaining high-quality nuclei were normalized and integrated using the standard Seurat SCTransform (v2) workflow based on the 3000 most variable genes while controlling for the number of genes per nucleus and the proportion of mitochondrial reads mapped. We then used default Harmony algorithm to correct for batch-effect differences. Clustering visualizations are projected via UMAP from 30 principal component analysis dimensions at a resolution of 0.2 and later customized using ggplot2.

#### Identification of marker features

Highly enriched genes in each cluster were determined by the ‘FindAllMarkers’ function, which identified the most enriched genes in a given cluster relative to all other clusters. Gene marker enrichment is defined as having a log_2_FC of ≥ 0.25 via MAST with per-cell gene number and mitochondrial reads as covariates. Cell type assignment was conducted by overlapping enriched snRNA-seq genes from each cluster with known markers of different liver cell types as defined in previous snRNA-seq papers. Differential expression analysis within each cell cluster between tissue status (FLC vs NML) using the ‘FindMarkers’ function in Seurat and significantly expressed genes are determined via MAST.

#### Inference of cell-cell communication patterns

Differences in cell-to-cell communication patterns between two biological conditions was conducted using CellChat (*36*) (v1.6.1). First, the Seurat object was subset into separate biological conditions, FLC or NML. The biological condition specific Seurat objects were converted into CellChat objects and run independently as described by the publicly available CellChat vignette. For either CellChat object, over expressed genes and interactions were identified using default parameters of the ‘identifyOverExpressedGenes’ and ‘identifyOverExpressedInteractions’ functions. Communication probabilities between cell types was computed using the function ‘computeCommunProb’ with the optional flags ‘type = “triMean”’ and ‘population.size = TRUÈ. Systems level analysis was conducted with the ‘identifyCommunicationPatterns’ function for ‘patterns = “incoming”’ or ‘patterns = “outgoing”. Functional and structural patterns were quantified using the ‘computeNetSimilarity’ function with ‘type = “functional”’ and ‘type = “structural”. After running CellChat on individual CellChat objects, FLC and NML CellChat objects were merged using ‘mergeCellChat’ and analyzed using default parameters. Heatmap, chord plot, and dotplot visualizations were generated using ‘netVisual_heatmap’, ‘netVisual_aggregatè, ‘netAnalysis_dot’ respectively.

## Functional studies

### si-SLC16A14 transfection in HepG2-DP+ clones

For the cell viability assay, HepG2-DP+ clones (H12 and H33) were seeded in a 96-well plate and transfected with 0.25-100 nM Silencer Select small interfering RNA (siRNA) (Negative Control: #1, SLC16A14: s90051) (Life Technologies) using Lipofectamine RNAiMAX transfection reagent (Life Technologies). Four days after the transfection, cell viability was measured using CellTiter-Glo 2 reagent (Promega). After the addition of the reagent to cell culture medium, luminescence was measured by CLARIOstar Plus microplate reader (BMG Labtech).

For RT-qPCR, HepG2-DP+ clones (H12 and H33) were seeded in a 6-well plate and transfected with 1-10 nM siRNA (Negative Control: #1, SLC16A14: s90051) using Lipofectamine RNAiMAX transfection reagent. Forty-eight hours after the transfection, total RNA was isolated from cells using the Qiagen RNeasy kit (Qiagen, Valencia CA). RT-qPCR was performed using GoTaq Probe 1-Step RT-PCR system (Promega) in a QuantStudio 7 Flex Real-Time PCR System (Applied Biosystems). The SLC16A14 mRNA level was measured with Taqman Gene Expression System (FAM-MGB) (Applied Biosystems) (Assay ID: SLC16A14: Hs00541300_m1, GAPDH: Hs02786624_g1). GAPDH mRNA levels were used as internal control and the data were analyzed using the ^ΔΔ^Ct method.

### siRNA-mediated knockdown of DNAJB1-PRKACA in a PDX-derived FLC cell line As described in Ma et al., 2024 (*11*)

#### Tissue slice preparation and drug treatments

The process for preparing tumor slices was carried out as previously described (*56*, *57*). In summary, tumor tissues were sliced into 400-micrometer sections using a Leica VT1200S vibratome (Leica Biosystems), employing HBSS as the slicing medium. These slices were further processed into 400-micrometer cuboids using the McIlwain tissue chopper (Ted Pella) as described in (*58*). These cuboids were immediately placed into 96-well ultralow-attachment plates (Corning) in Williams’ medium supplementing with 12 mM nicotinamide, 150 nM ascorbic acid, 2.25 mg/mL sodium bicarbonate, 20 mM HEPES, an additional 50 mg/mL glucose, 1 mM sodium pyruvate, 2 mM L-glutamine, 1% (v/v) ITS, 20 ng/mL EGF, 40 IU/mL penicillin, and 40 ug/mL streptomycin. The RealTime Glo reagent (Promega) was added to the incubation media as per the guidelines provided by the manufacturer. The baseline cell viability in the cuboids was determined after 24 hours using RealTime Glo bioluminescence, measured with a Synergy H4 instrument (Biotek). The cuboids were subjected to either DMSO as a control or various experimental drugs as indicated. Measurements of the overall viability of the tumor tissues were taken daily for up to 7 days following the treatment.

#### Gene knockdown experiments in tissue slices

The preparation of tissue slices followed the method as described above. We obtained small interfering RNA (siRNA) targeting SDS, or non-targeting control from Dharmacon (Thermo Fisher Scientific). To evaluate changes in aggregate viability, we performed siRNA transfections in 96-well plates using Lipofectamine RNAiMax (provided by Invitrogen), following the instructions of the manufacturer. Each transfection process involved a minimum of two wells, with each well containing 4 to 6 3D cuboids. We monitored the viability changes using the Synergy H4 instrument from Biotek. To assess the overall changes in the viability of the tumor tissues, we carried measurements for a period of up to 7 days after the treatment.

#### PDX-derived FLC cell line knockdown

The PDX-derived FLC primary cell line (FLX1) was transfected at two timepoints (day 1 and day 4) during a six-day trial with 500nM of siRNA negative control (ThermoFisher Scientific; Cat No.—12935200) or siRNA against SLC16A14 (ThermoFisher Scientific; Cat No.— 1299001) using lipofectamine 3000 (ThermoFisher Scientific; Cat No.—L3000015) in a 24-well plate format (n = 2 per condition) for assessment of knockdown efficiency and 96-well plate format (n =3 per condition) for assessment of cell viability. On day 6, RNA was isolated from harvested cells using the Norgen Total RNA purification kit (Norgen Biotek Corp; Cat No.— 37500) and used for cDNA generation (BIORAD T100 Thermal Cycler) which served as input for qPCR (BIORAD C1000 Touch Thermal Cycler). SLC16A14 quantification (Target probe from ThermoFisher Scientific; Cat No.—4331182; Assay ID—Hs00541300_m1) was normalized to RPS9 (Target probe from ThermoFisher Scientific; Cat No.—4351370; Assay ID—Hs02339424_g1). On day 6, the Cell-Titer Glo kit (Cat No.—G7570) was applied to cells, and chemiluminescence was assessed on the Biotek Synergy 2 with the following settings: read mode—area scan, read height—1mm, optics position—top, gain—135, emission—hole, and integration time—50ms.

#### Secreted protein assessment

Conditioned media from FLC tumor slices and normal hepatocytes were collected after 4 days in culture and sent to NomicBio (Montreal, Canada) for nELISA-based secreted protein analysis as described previously (*59*, *60*).

## Supplemental figure legends

**Fig. S1.**
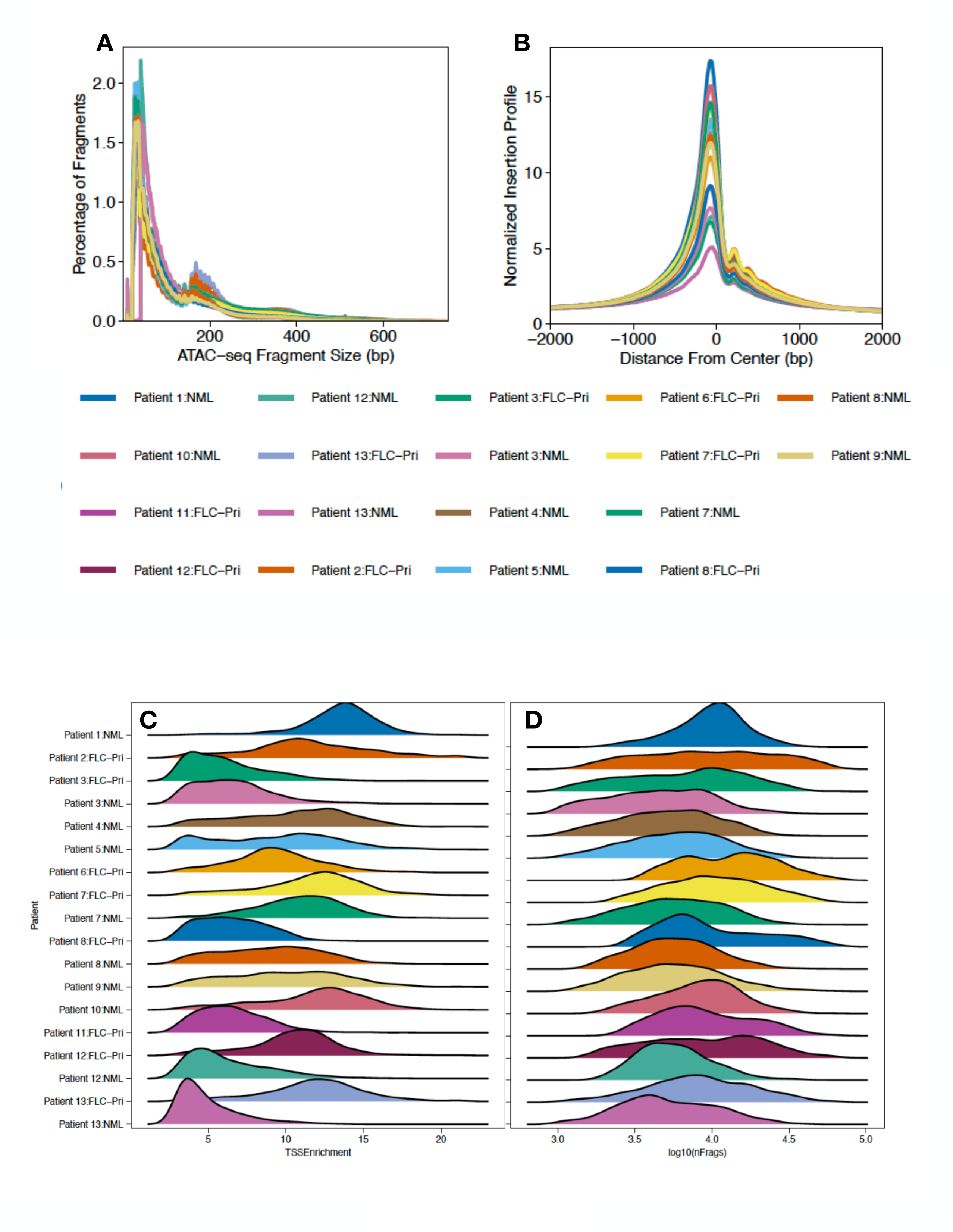
Sample statistics and quality control metrics for snATAC-seq dataset. (**A**) Fragment size distribution. (B) TSS insertion profiles centered at all annotated TSS loci. (C and D) Ridge plots depicting quality of each snATAC-seq dataset. The distribution of (left) TSS enrichment and number of unique fragments (right) using a log10 scale. FLC-Pri, fibrolamellar carcinoma-primary tumor; FLC-Met, fibrolamellar carcinoma-metastatic tumor; FLC-Recurr, fibrolamellar-recurrent tumor; NML, non-malignant liver.

**Fig. S2.**
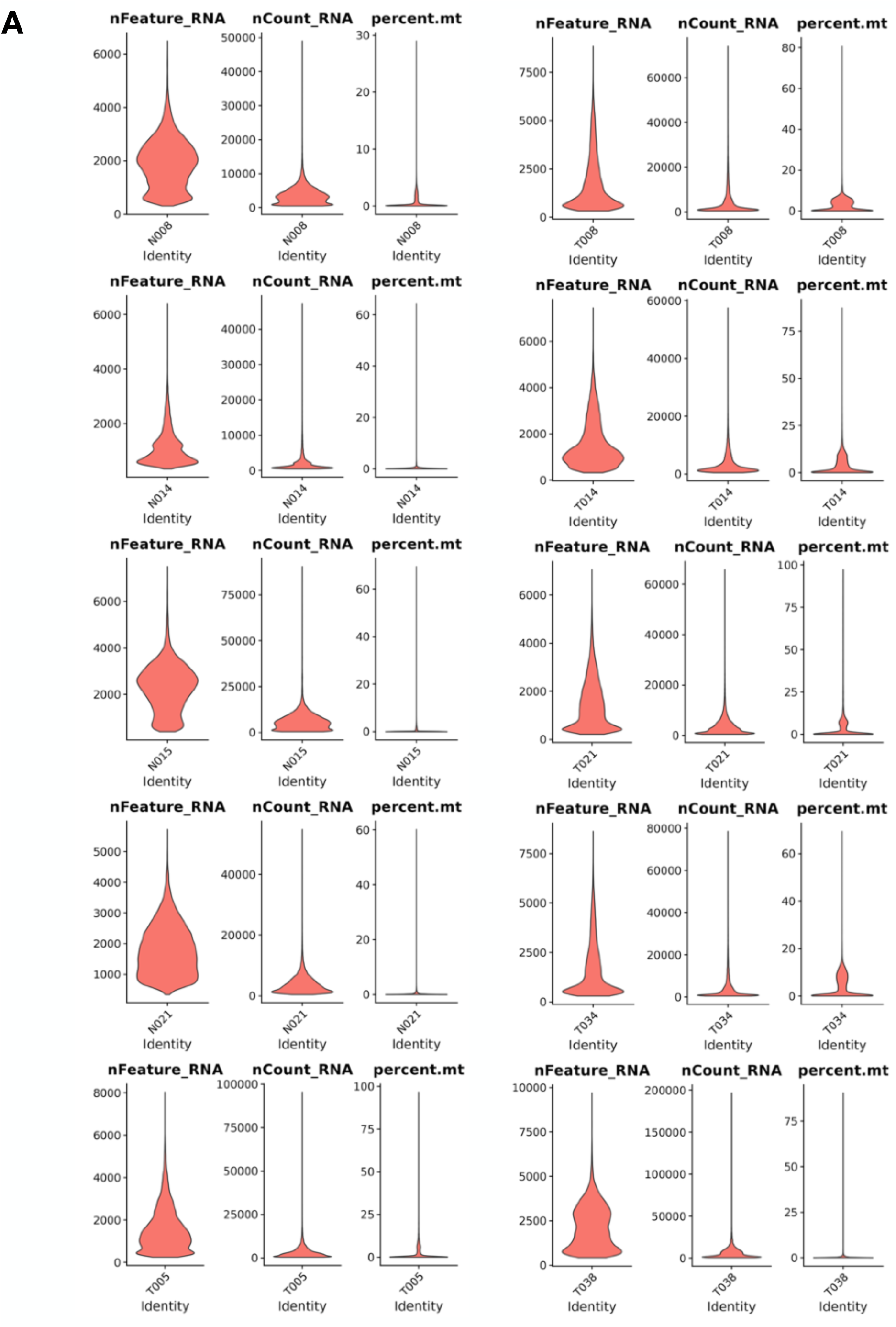
Sample statistics and quality control metrics for snRNA-seq dataset. (A) Violin plots depicting the sample statistics of the (nFeature_RNA, left) number of unique genes per cell, (nCount_RNA, middle) number of RNA molecules detected by unique molecular identifier, and (percent.mt, right) percent mitochondrial RNA, captured in each sample. (B) Table containing the number of cells from each sample before and after quality control filtering (nFeature > 1000 & percent.mt < 20%).

**Fig. S3.**
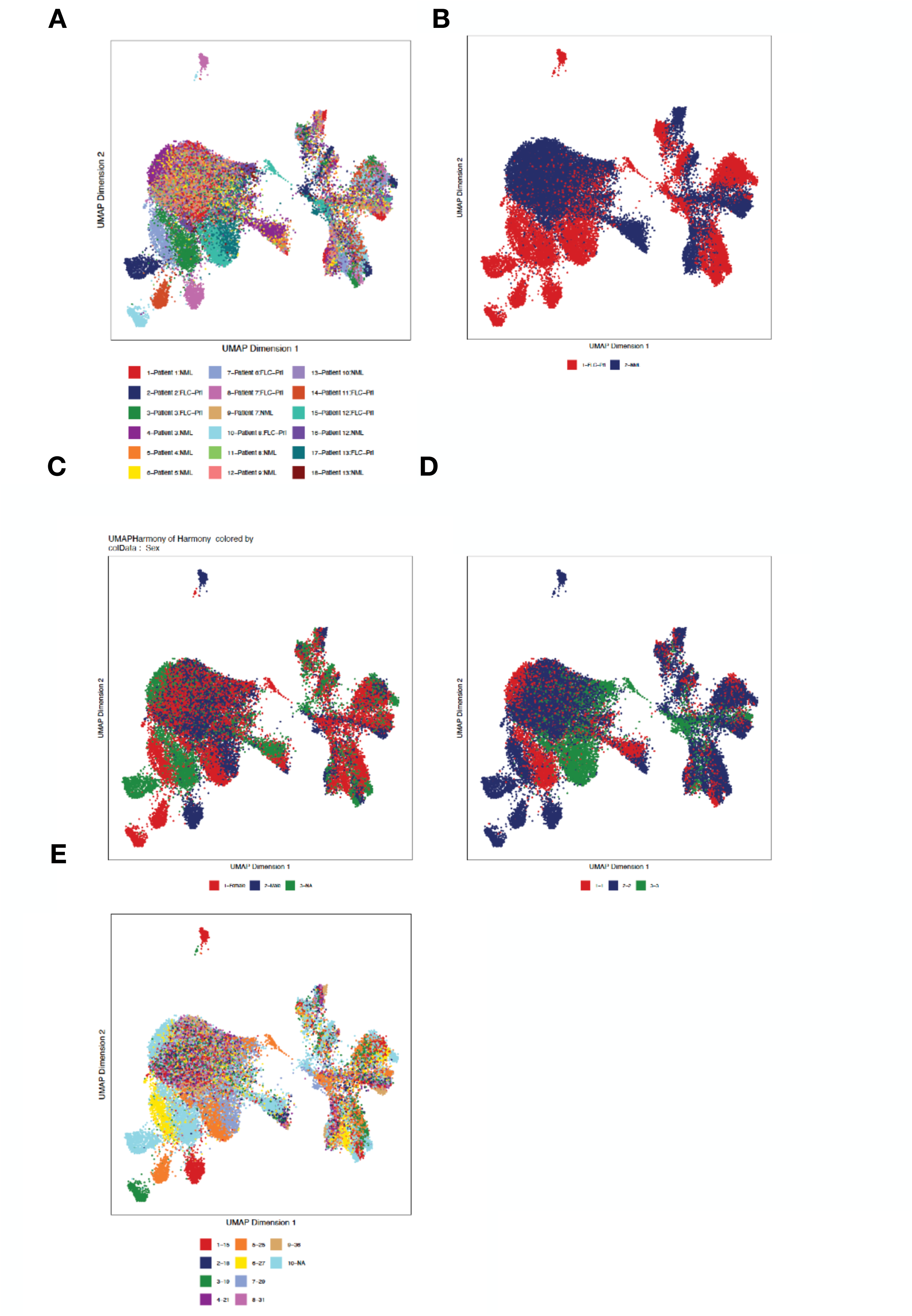
snATAC-seq UMAP overlays of known metadata. (A-E) UMAP embedding overlayed with (A) patient and tissue status information, (B) tissue status, (C) patient sex information, (D) sequencing batch, and (E) age. FLC-Pri, fibrolamellar carcinoma-primary tumor; FLC-Met, fibrolamellar carcinoma-metastatic tumor; FLC-Recurr, fibrolamellar-recurrent tumor; NML, non-malignant liver.

**Fig. S4.**
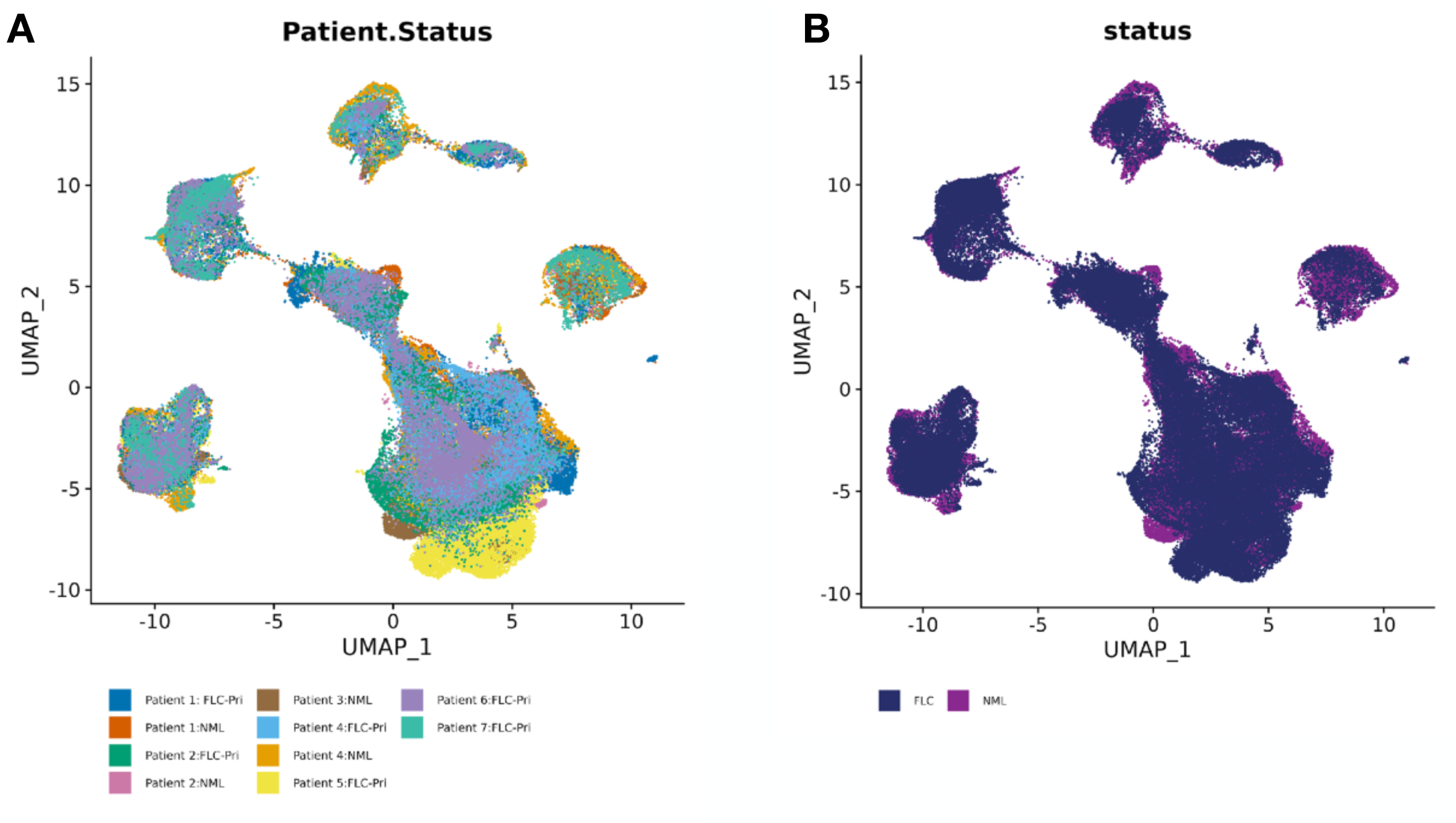
snRNA-seq UMAP overlays of known metadata. (A and B) UMAP embedding overlayed with (A) Patient and tissue status information and (B) tissue status.

**Fig. S5.**
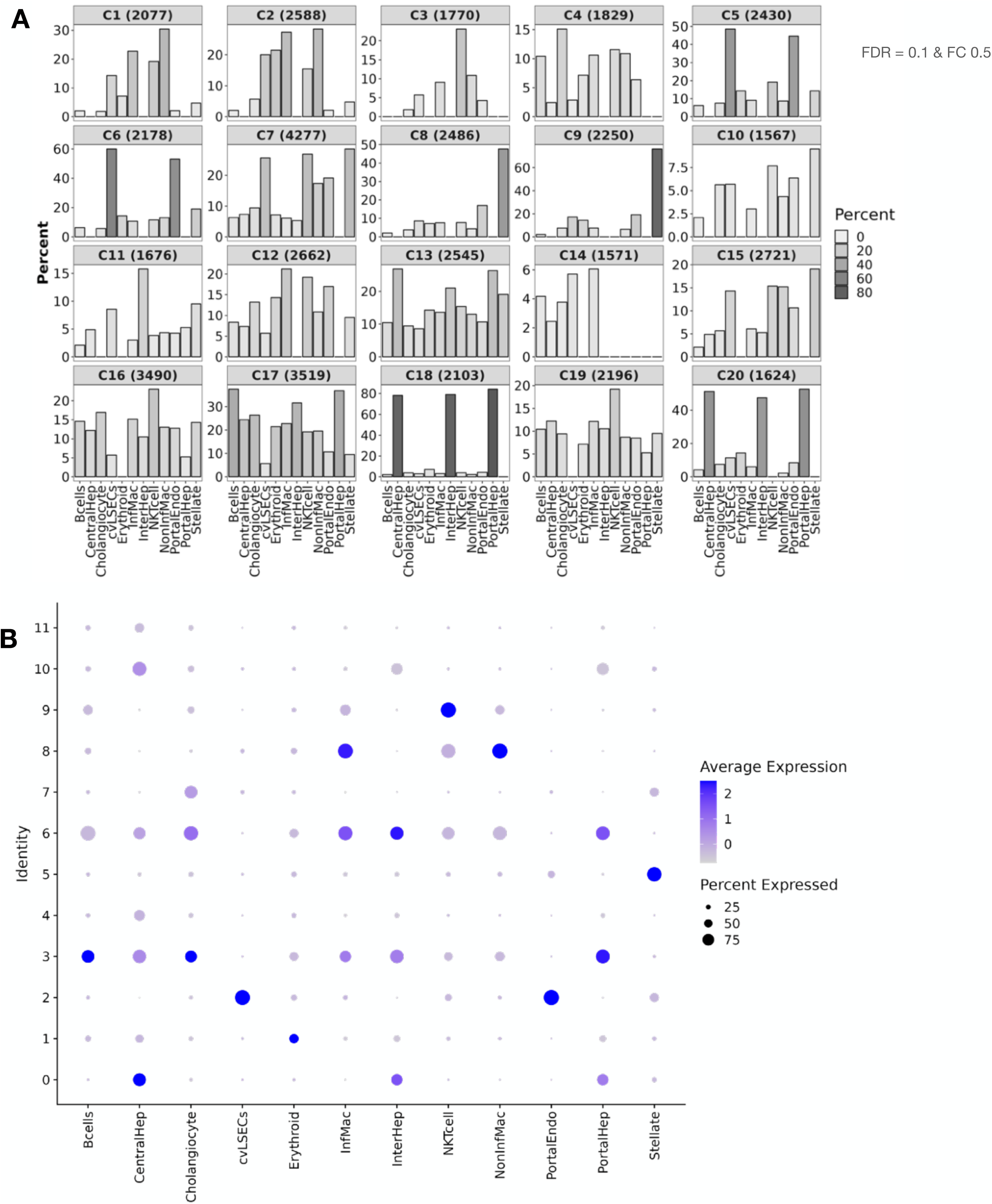
A priori cell type assignment strategy reveals cluster identities in snATAC-seq and snRNA-seq data. (A) snATAC-seq: overlap analysis of marker genes of each cluster with bona fide liver cell markers. (B) snRNA-seq: module scores depicting gene expression of different liver cell markers within each cluster.

**Fig. S6.**
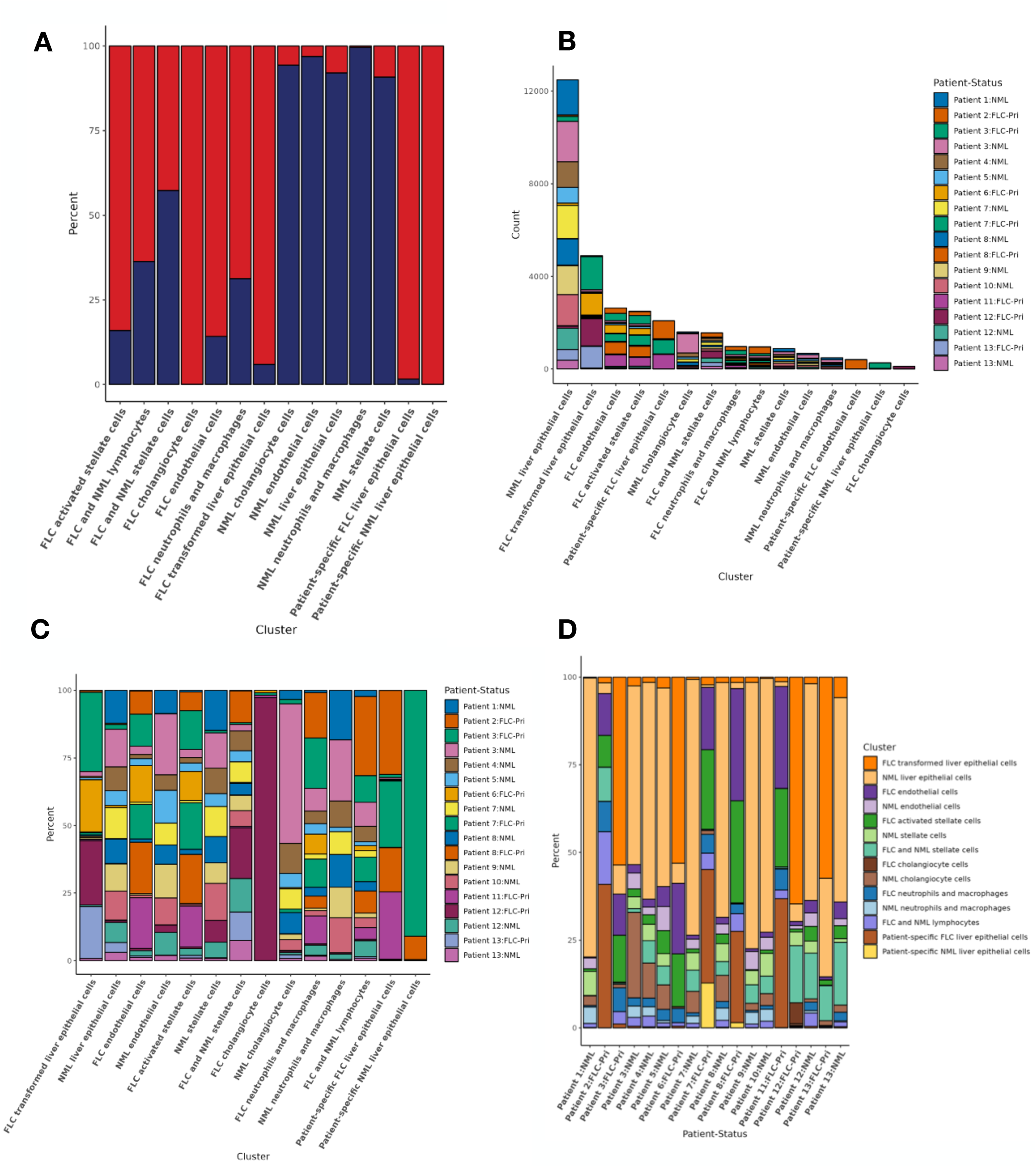
snATAC-seq cell proportions. (A) Stacked bar plot depicting tissue status proportion within each cell type. (B) Histogram of total number of cells within each cluster and contribution from each patient sample. (C) Stacked bar plot illustrating the cell type breakdown by patient sample. (D) Stacked bar plot depicting cell type contribution by each patient sample. FLC-Pri, fibrolamellar carcinoma-primary tumor; FLC-Met, fibrolamellar carcinoma-metastatic tumor; FLC-Recurr, fibrolamellar-recurrent tumor; NML, non-malignant liver.

**Fig. S7.**
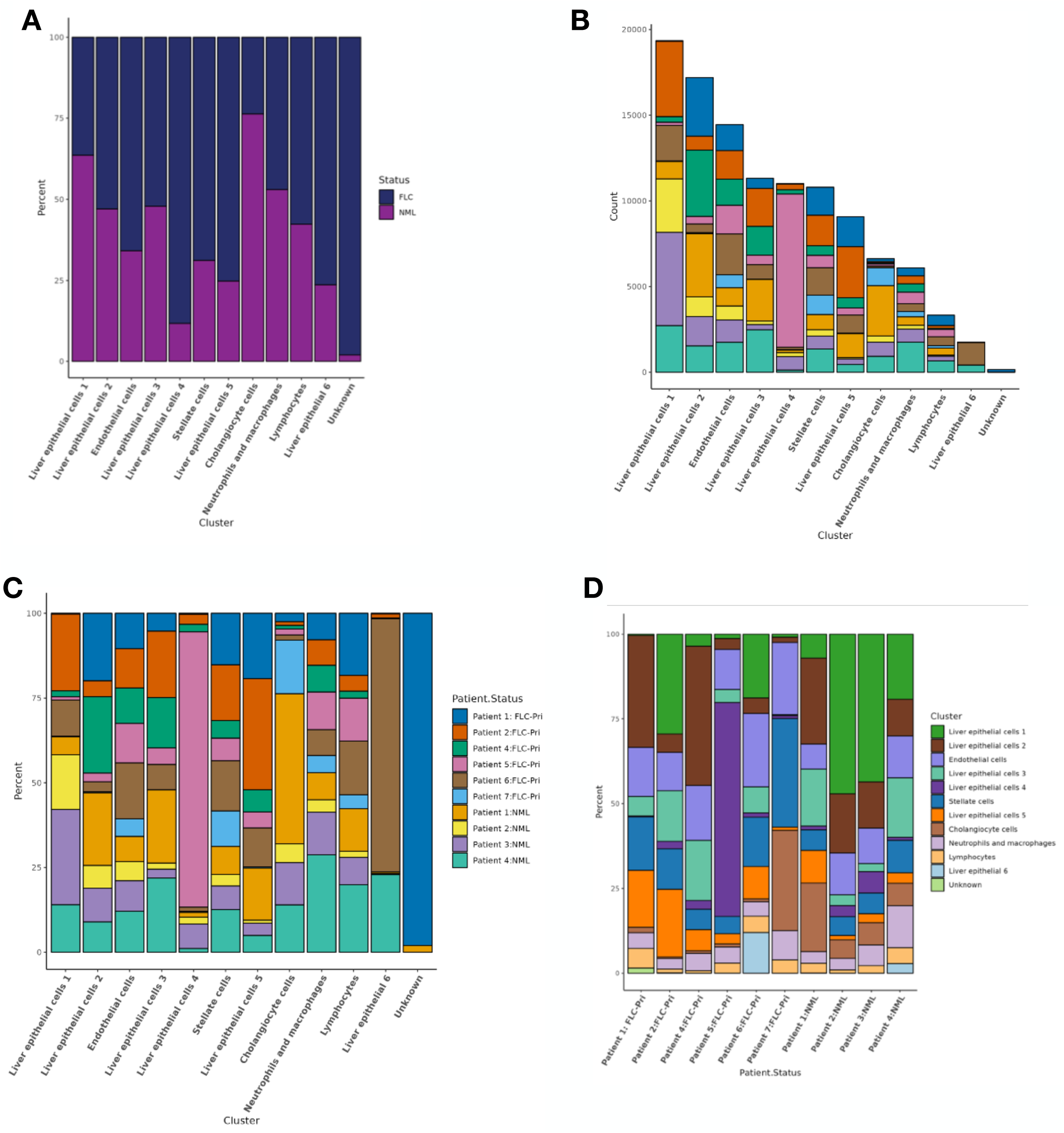
snRNA-seq cell proportions. (A) Stacked bar plot depicting tissue status proportion within each cell type. (B) Histogram of total number of cells within each cluster and contribution from each patient sample. (C) Stacked bar plot illustrating the cell type breakdown by patient sample. (D) Stacked bar plot depicting cell type contribution by each patient sample. FLC, fibrolamellar carcinoma tumor; NML, non-malignant liver.

**Fig. S8.**
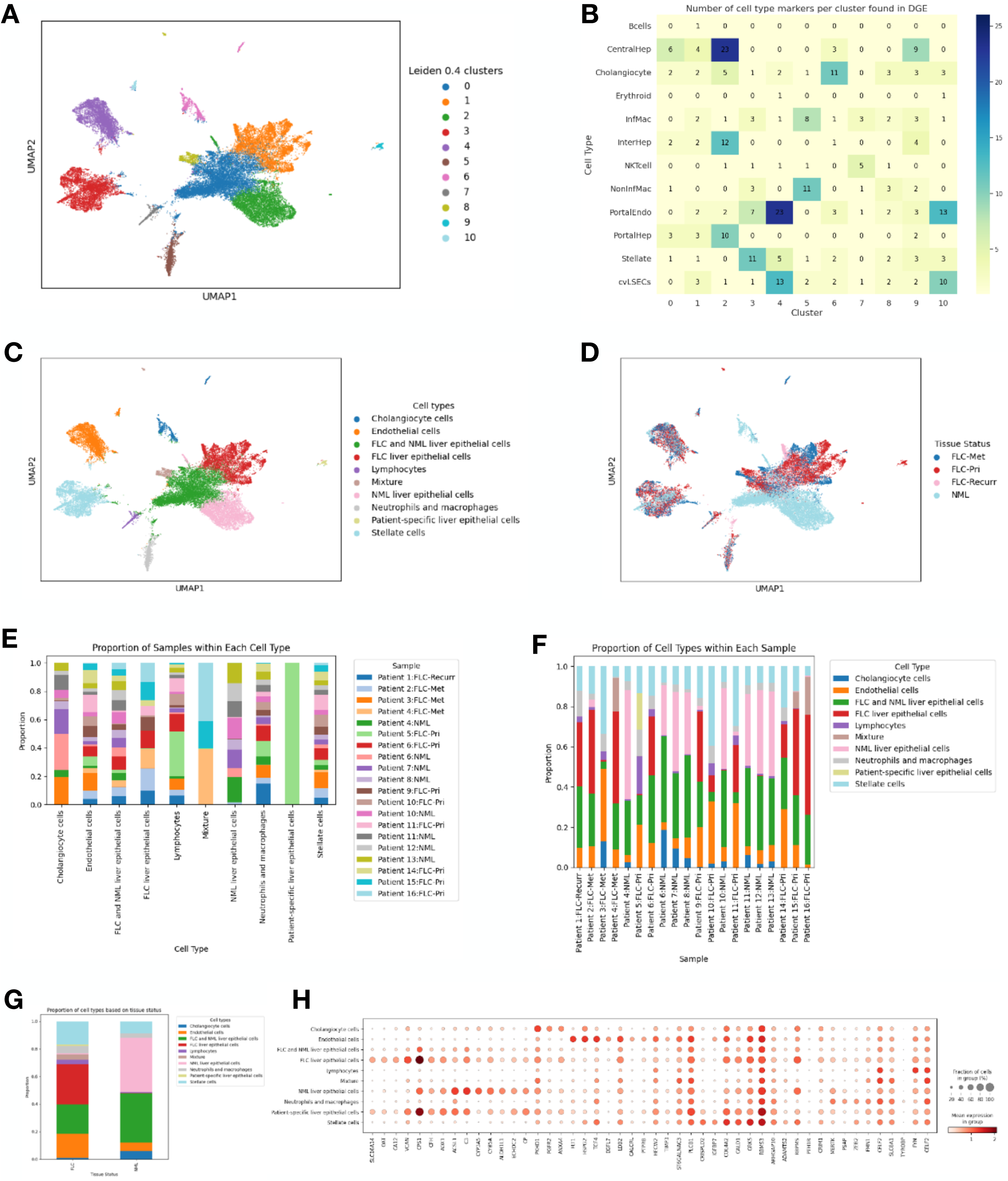
Orthogonal snATAC-seq analysis by SnapATAC2 corroborates ArchR analysis. (A) Unannotated UMAP overlay of snATAC-seq data using SnapATAC2. (B) Overlap analysis of marker genes of each cluster with bona fide liver cell markers. (C) Annotated UMAP overlay of snATAC-seq data using SnapATAC2. (D) UMAP overlay of patient and tissue status information. (E) Stacked bar plot illustrating the cell type breakdown by patient sample. (F) Stacked bar plot depicting cell type contribution by each patient sample. (G) Stacked bar plot showing proportions of cell types depicted in FLC (left) and NML (right). (H) Dotplot depicting gene activity scores of FLC genes in each cell type. Genes are labeled at the bottom of the dotplot. FLC-Pri, fibrolamellar carcinoma-primary tumor; FLC-Met, fibrolamellar carcinoma-metastatic tumor; FLC-Recurr, fibrolamellar-recurrent tumor; NML, non-malignant liver.

**Fig. S9.**
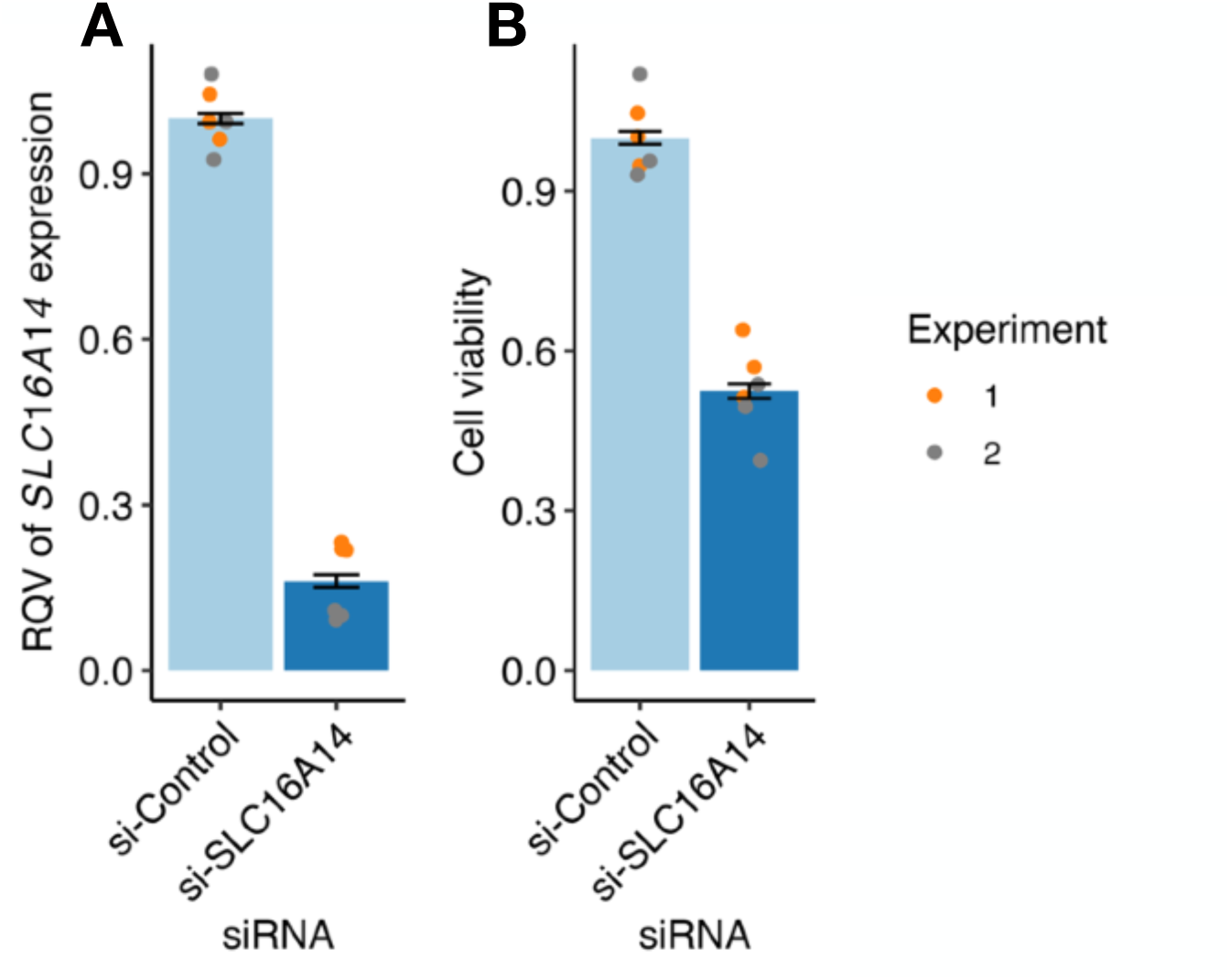
Knockdown SLC16A14 in PDX-derived FLC cell line leads to reduced cell viability. (A) Relative *SLC16A14* expression in a PDX-derived FLC cell line (FLX1, n= 2 independent experiments, 3 technical replicates each, for a total of 6 replicates) after treatment with si-SLC16A14 (500nM). (B) Cell viability of PDX-derived FLC cells after si-SLC16A14 treatment (500 uM).

**Fig. S10.**
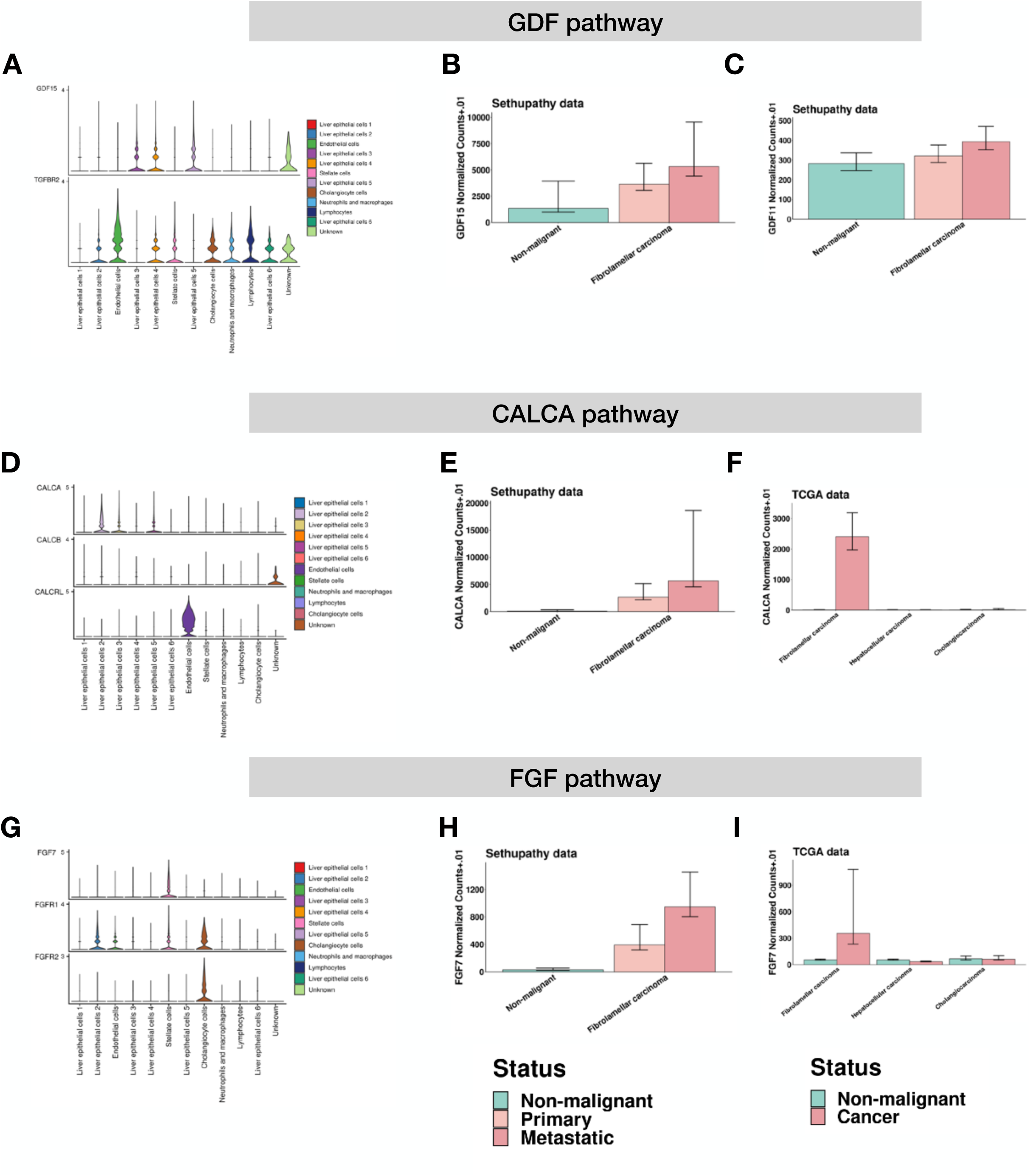
CellChat details newly activated ligand-receptor pairs that are unique to FLC. (A, D, and G) Violin plots depicting expression of ligands and receptors for (A) GDF, (D) CALCA, and (D) FGF pathways. (B, C, E, and H) Bar plots depicting the expression of (A) GDF15, (C) GDF11, (E) CALCA, and (H) FGF7 in NML (left), FLC-Pri (center), and FLC-Met (right). (F and I) Bar plots depicting the expression of (F) CALCA and (I) FGF7 in FLC (left), HCC (middle) and CCA (right). FLC-Pri, fibrolamellar carcinoma-primary tumor; FLC-Met, fibrolamellar carcinoma-metastatic tumor; NML, non-malignant liver; CCA, cholangiocarcinoma.

**Fig. S11.**
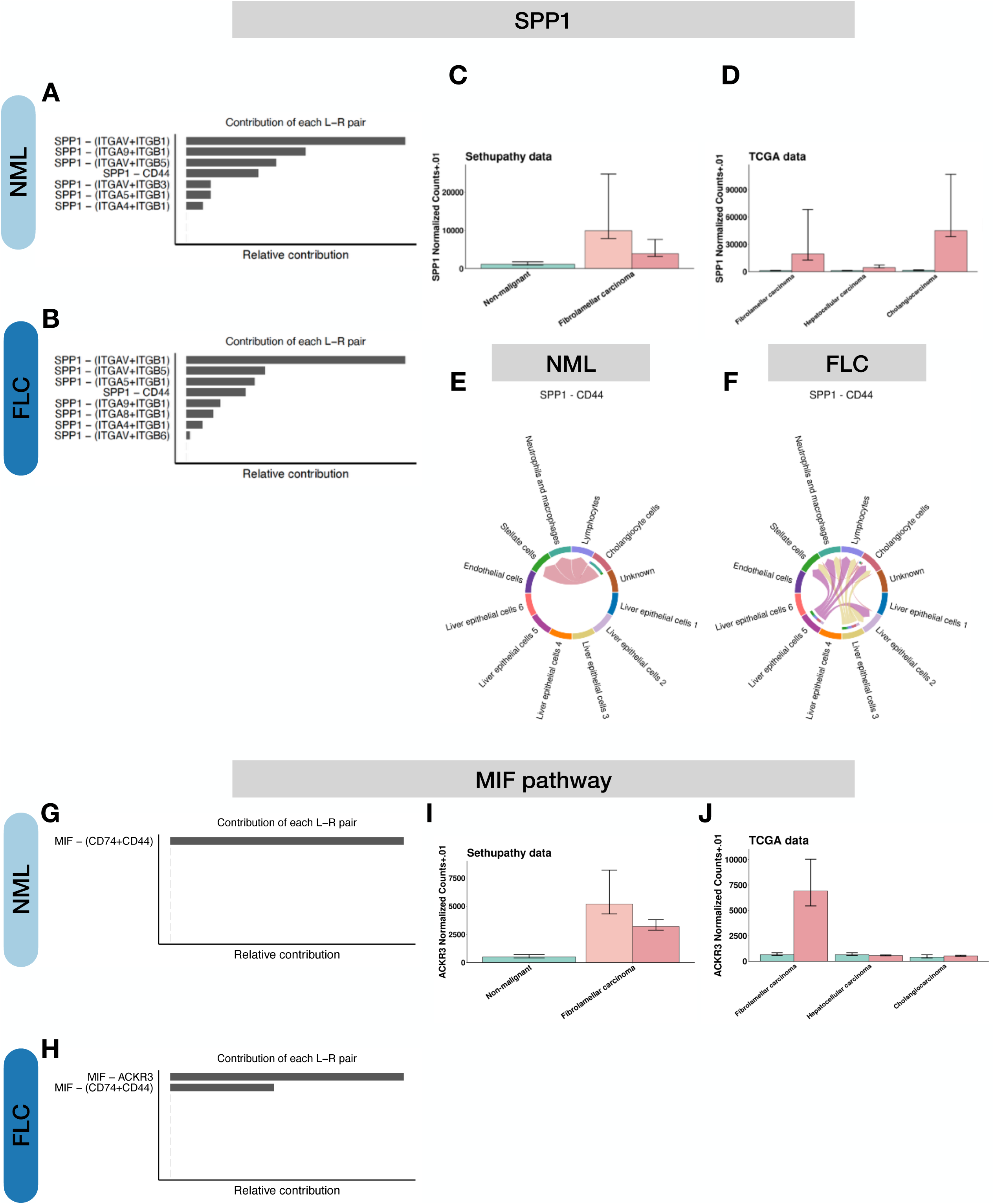
CellChat details rewired pathways in FLC relative to NML. (A,B, G-H) Relative contributions of ligand-receptor pairs for SPP1 (A-F) and MIF (G-J) pathways in NML (A, G; top) or FLC (B, H; bottom). (C and I) Bar plots depicting the expression of SPP1 (top; C) or ACKR3 (bottom; I) in NML (left), FLC-Pri (center), and FLC-Met (right). (D and J) Bar plots depicting the expression of SPP1 (top; D) or ACKR3 (bottom; J) in FLC (left), HCC (middle) and CCA (right).

**Fig. S12.**
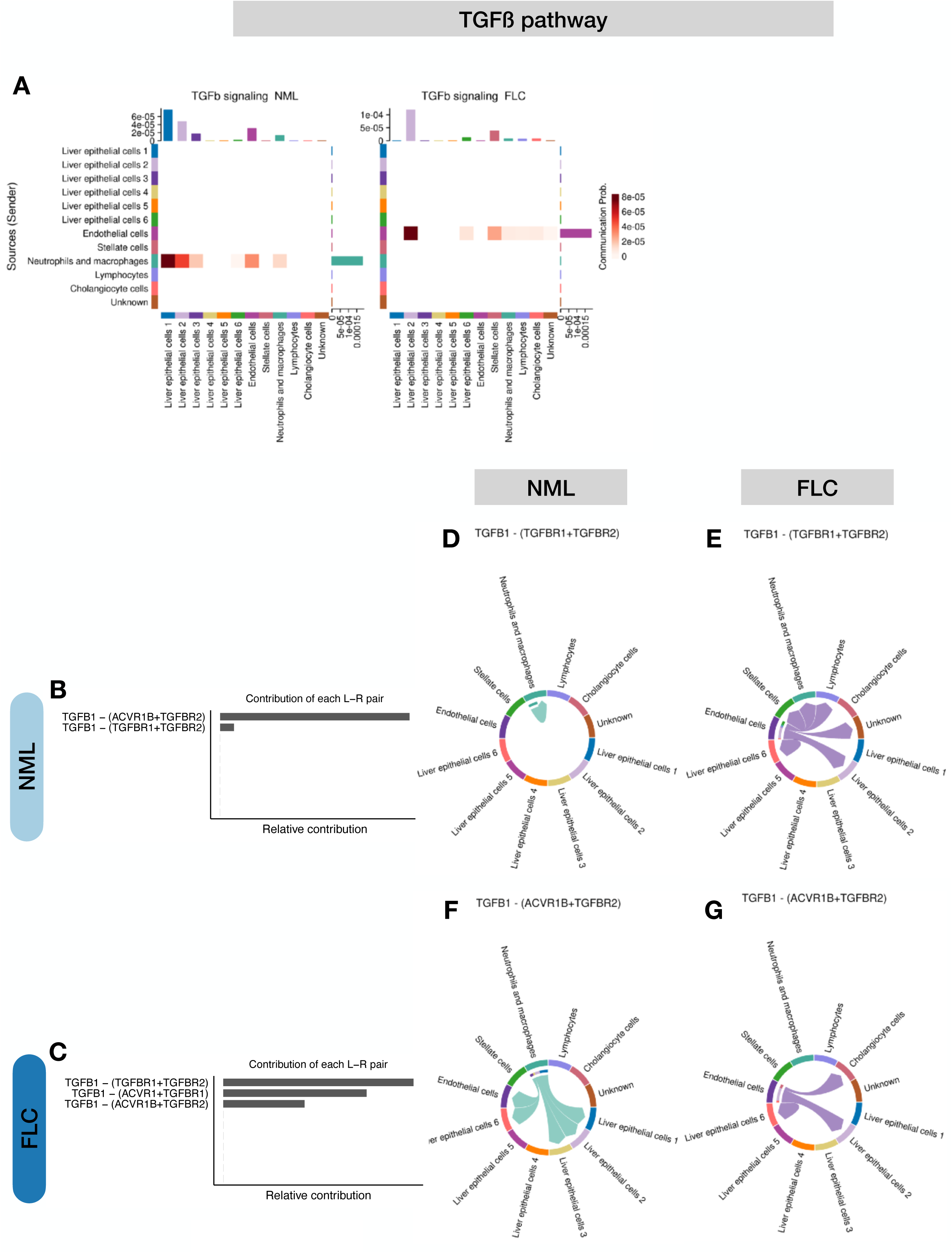
CellChat details TGFB rewiring. (A) Heatmap depicting TGFB signaling pathways in FLC. Source (sender) cells are labeled on the left, and the receiver cell type(s) is/are labeled on the bottom. The right bar graph depicts sum of contribution from sources. The bar graphs on top depict the sum of column values for receptors. (B and C) Relative contributions of ligand-receptor pairs for TGFß pathway in NML (B; top) and FLC (C; bottom). (D and E) Chord plots depicting TGFβ1-(TGFβR1+TGFβR2) wiring in NML (D) and FLC (E). (F and G) Chord plots depicting TGFβ1-(ACVR1B+TGFβR2) wiring in NML (F) and FLC (G). FLC-Pri, fibrolamellar carcinoma-primary tumor; FLC-Met, fibrolamellar carcinoma-metastatic tumor; NML, non-malignant liver; CCA, cholangiocarcinoma.

## Acknowledgements

This work was supported by different Fibrolamellar Cancer Foundation Research Grants (awarded separately to PS and TSG) as well as by funding from the Sandra Atlas Bass Endowment Grant (awarded to PS), the Tisch Families Faculty Development Fund and CHMC Surgical Foundation, Inc. (awarded to KV), the National Cancer Institute (R01CA273081, awarded to TSG), and the Cornell Provost’s Diversity Fellowship for Advanced Doctoral Students (awarded to ARF). Department of Defense (CA180067); Fibrolamellar Cancer Foundation. LKD is a recipient of the Cancer Research Institute/Fibrolamellar Cancer Foundation Postdoctoral Fellowship (CRI Award 4093).

---

**Table S1.**

| **Sample** | **Cells** | **After filtering**<br>**(minTSS >= 3 &**<br>**nFrag >= 1000)** | **Doublet Removal** |
| --- | --- | --- | --- |
| **Patient 1:NML** | 1965 | 1944 | 1907 |
| **Patient 2:FLC-Pri** | 2070 | 1929 | 1892 |
| **Patient 3:FLC-Pri** | 3908 | 2709 | 2673 |
| **Patient 3:NML** | 5360 | 3471 | 3408 |
| **Patient 4:NML** | 2070 | 1819 | 1786 |
| **Patient 5:NML** | 1798 | 1214 | 1200 |
| **Patient 6:FLC-Pri** | 1969 | 1809 | 1777 |
| **Patient 7:FLC-Pri** | 1970 | 1928 | 1891 |
| **Patient 7:NML** | 2072 | 1998 | 1959 |
| **Patient 8:FLC-Pri** | 1973 | 1594 | 1569 |
| **Patient 8:NML** | 1972 | 1714 | 1685 |
| **Patient 9:NML** | 1979 | 1679 | 1651 |
| **Patient 10:NML** | 1978 | 1886 | 1851 |
| **Patient 11:FLC-Pri** | 1975 | 1722 | 1693 |
| **Patient 12:FLC-Pri** | 1960 | 1839 | 1806 |
| **Patient 12:NML** | 1970 | 1514 | 1492 |
| **Patient 13:FLC-Pri** | 1824 | 1661 | 1634 |
| **Patient 13:NML** | 1890 | 656 | 652 |
| **Total** | **40703** | **33086** | **32526** |

**Table S2.**

| **Sample** | **Cells** | **After doublet removal** | **After filtering (nFeature_RNA > 1000)** |
| --- | --- | --- | --- |
| **N008** | 21713 | 18130 | 14503 |
| **N014** | 16844 | 14662 | 6625 |
| **N015** | 17425 | 14426 | 12491 |
| **N021** | 20046 | 17637 | 14132 |
| **T005** | 25734 | 22116 | 14172 |
| **T008** | 23090 | 18612 | 10388 |
| **T014** | 27915 | 23717 | 14875 |
| **T021** | 19214 | 16820 | 9412 |
| **T034** | 24953 | 21607 | 10994 |
| **T038** | 5026 | 4676 | 3541 |
| **Total** | **201960** | **172403** | **111133** |

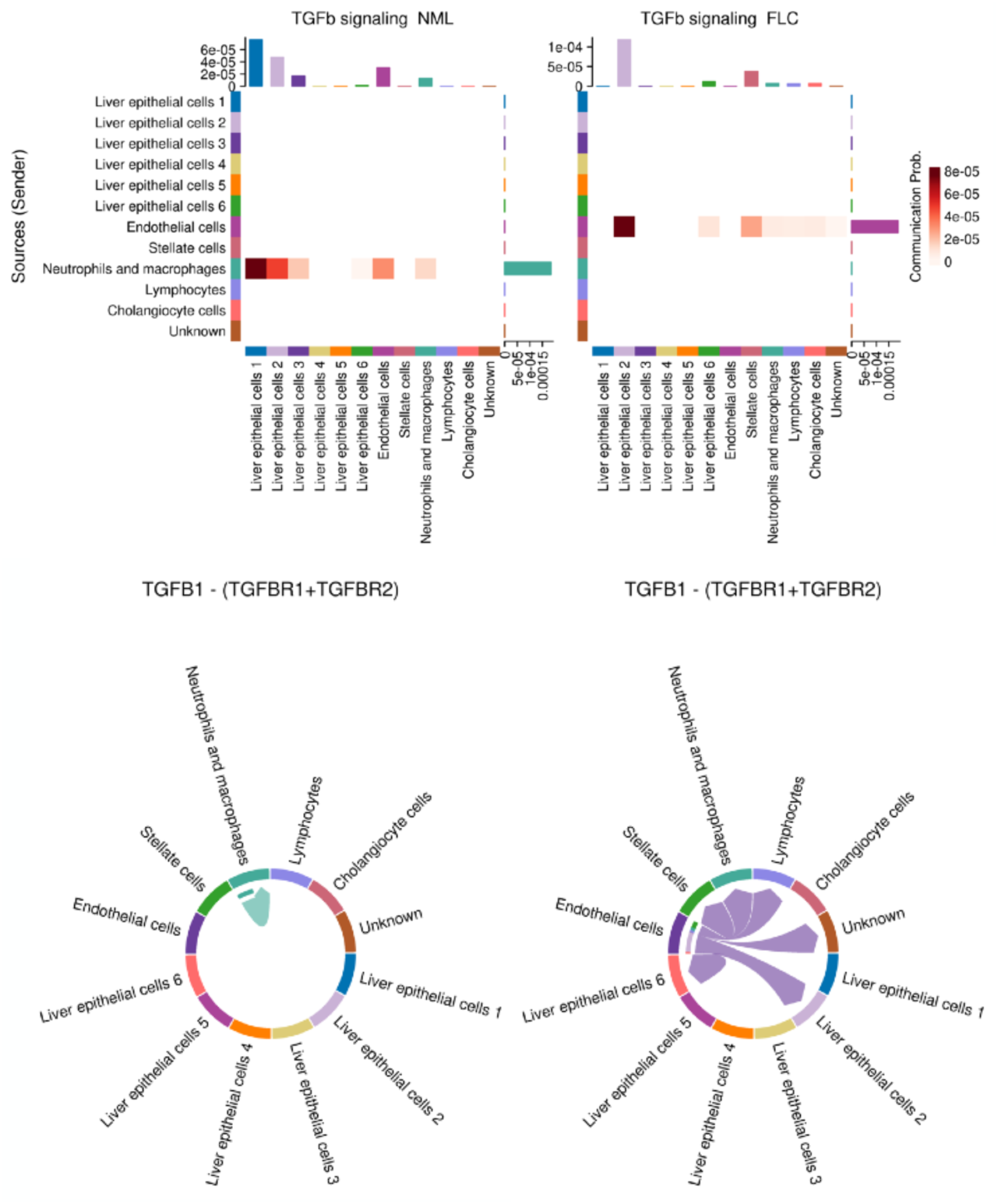

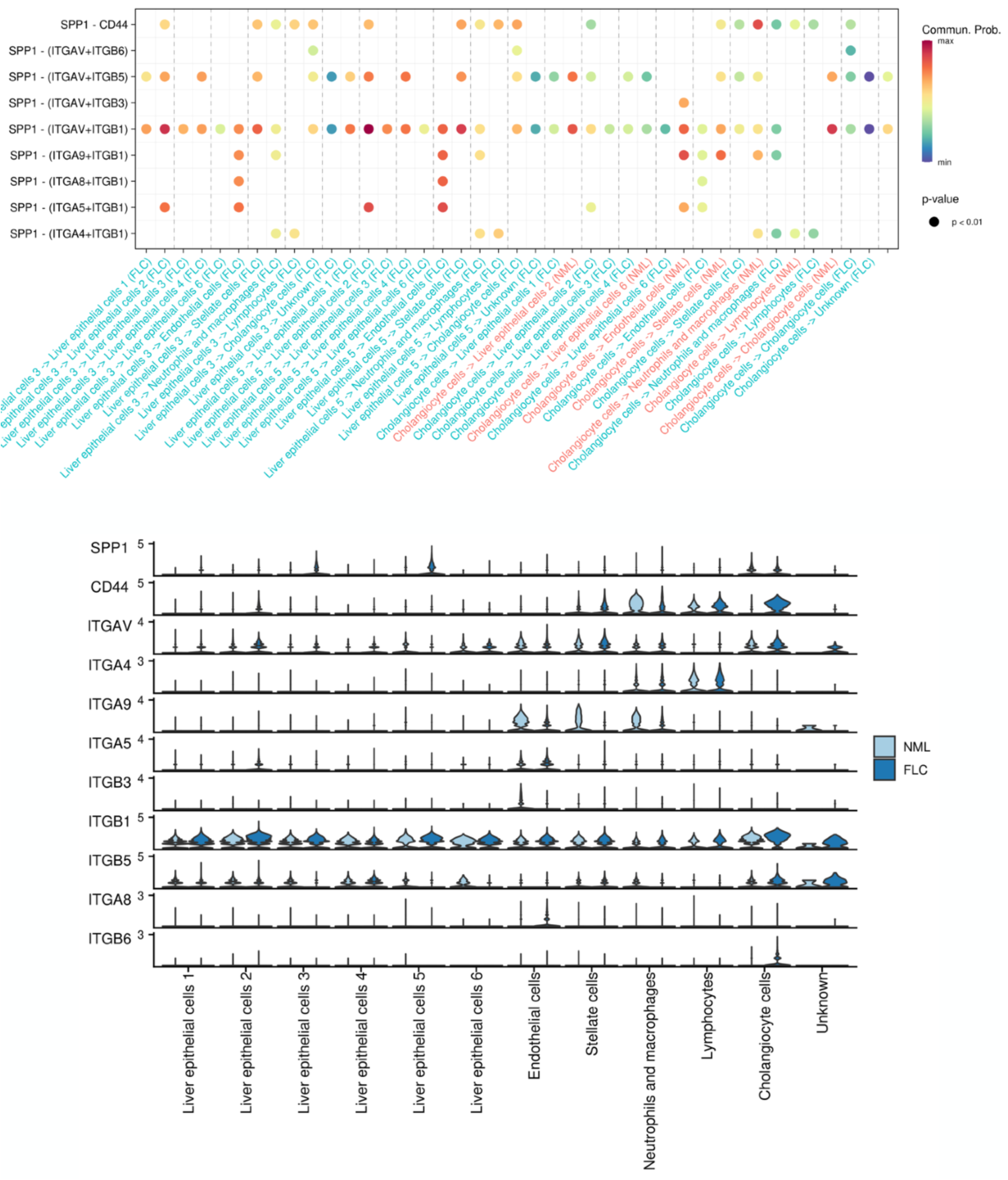

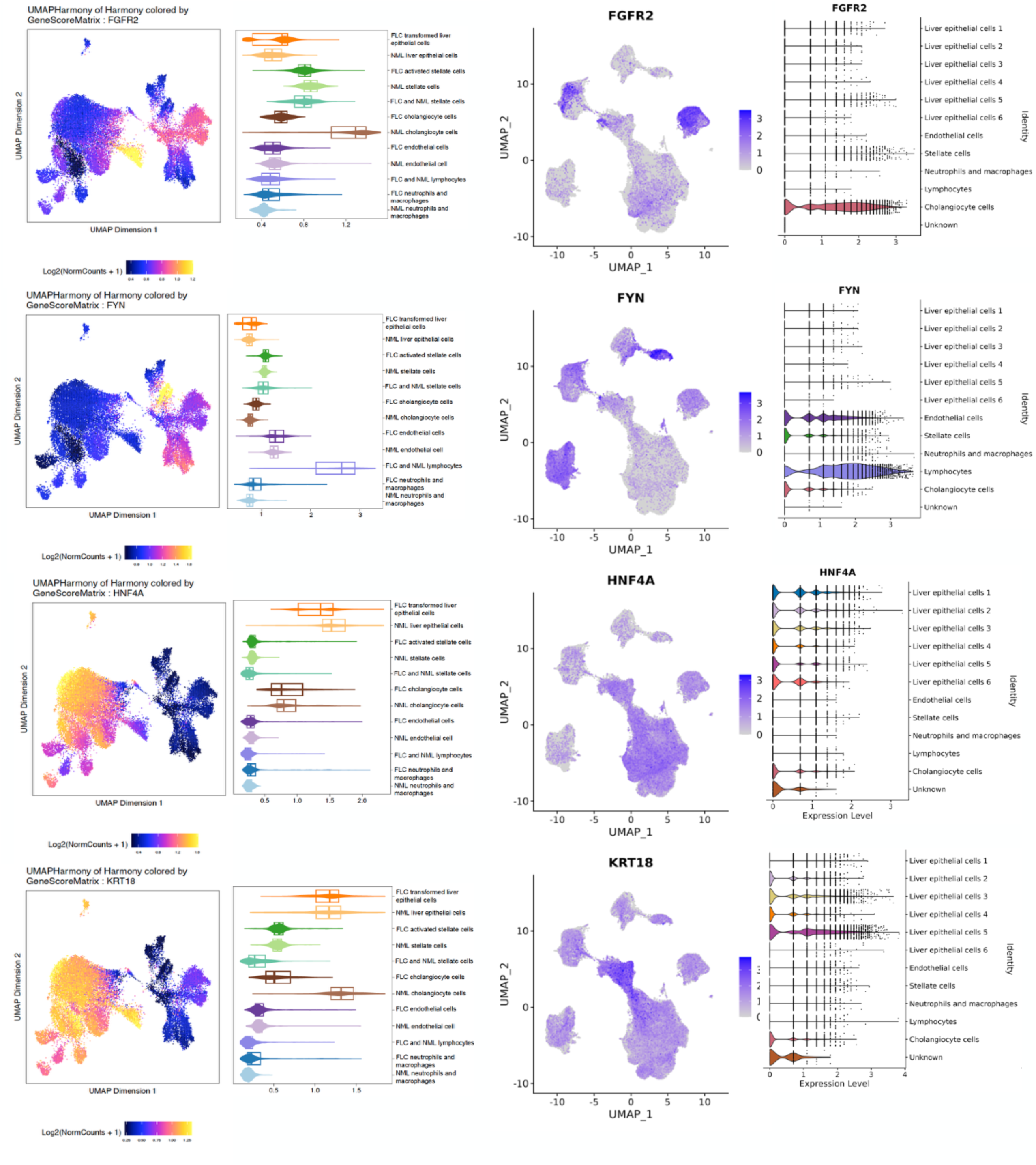

